# A unifying model to predict variable drug response for personalised medicine

**DOI:** 10.1101/2020.03.02.967554

**Authors:** Maaike van der Lee, William G. Allard, Rolf H.A.M Vossen, Renée F. Baak-Pablo, Roberta Menafra, Birgit A.L.M. Deiman, Maarten J. Deenen, Patrick Neven, Inger Johansson, Stefano Gastaldello, Magnus Ingelman-Sundberg, Henk-Jan Guchelaar, Jesse J. Swen, Seyed Yahya Anvar

## Abstract

Pharmacogenomics is a key component of personalized medicine. It promises a safer and more effective drug treatment by individualizing the choice of drug and dose based on an individual’s genetic profile^1,2^. The majority of commonly prescribed drugs are metabolized by a small set of Cytochrome P450 (CYP) enzymes^3^. In clinical practice, genetic biomarkers are being used to categorize patients into predefined *-alleles to predict CYP450 enzyme activity and adjust drug dosages accordingly. Yet, this approach has important limitations as it leaves a large part of variability in drug response unexplained^4,5^. Here, we present a novel approach and introduce a continuous scale (instead of categorical) assignments to predict metabolic enzyme activity. The proposed strategy uses full gene sequencing data, a neural network model and CYP2D6 mediated tamoxifen metabolism from a prospective study of 561 breast cancer patients. The model explains 79% of the interindividual variability in CYP2D6 activity compared to 54% with the conventional approach. It is capable of assigning accurate enzyme activity to alleles containing previously uncharacterized combinations of variants and were replicated in an independent cohort of tamoxifen treated patients, a cohort of Venlafaxine users as well as in vitro functional assays using HEK cells. These results demonstrate the advantage of a continuous scale and a completely phased genotype for prediction of CYP450 enzyme activity and thereby enables more accurate prediction of individual drug response.

---

The cytochrome P450 isoenzyme 2D6, encoded by the polymorphic *CYP2D6*^6^, is involved in the metabolism of 25-30% of commonly prescribed drugs^7^. Genetic variants in the *CYP2D6* gene, such as SNVs (single nucleotide variants), CNVs (copy-number variants) and structural rearrangements^6,8,9^, may lead to differential CYP2D6 activity and thereby to altered drug response ^10,11^.

To translate *CYP2D6* variants into clinically actionable guidelines, they are assigned to standard haplotypes and predicted phenotypes. Haplotype assignment is performed based on *-allele nomenclature, catalogued by the Pharmacogene Variation Consortium (PharmVar), where each *-allele describes a predefined combination of variants^12,13^. Subsequently, the gene activity score (GAS) system assigns a score to each allele, with 0 for no activity, 0.5 for decreased, 1 for normal and 2 for increased activity^14^. Predicted phenotypes are assigned based on the combination of the two inferred allele activities and are summarized into 4 different CYP2D6 metabolizer categories^13,15^: poor metabolizer (PM), intermediate metabolizer (IM), normal metabolizer (NM) and ultra-rapid metabolizer (UM). However, 6 to 22-fold unexplained intra-category variability in enzyme activity and considerable overlap in activity between phenotypes remains^5^. Moreover, a recent twin study has shown that while 91% of CYP2D6 metabolism is hereditary, GAS based inferred phenotypes only explained 39% of variability in CYP2D6 enzyme activity^4^. Similar trends in missing heritability have been shown for other genes involved in CYP450 mediated drug metabolism^16,17^. This is partly due to rare genetic variants that are not catalogued in the current *-allele nomenclature^18^. A major limitation of the current methodology is the loss of a considerable amount of information in the categorization. Therefore, a continuous phenotype prediction rather than a categorical model is likely to improve the prediction of CYP2D6 enzyme activity. A convolutional neural network is highly suitable for this type of phenotype prediction from genetic data^19,20^. While previous approaches of deep learning in pharmacogenomics were aimed at automated *-allele assignment or to predicting the residual activity of conventional *-alleles^21,22^, we propose a novel methodology to predict CYP2D6 enzyme function on a continuous scale using full gene sequencing data and a neural network, omitting *-alleles completely.

## Conventional categorical phenotype predictions

To study the explained variability of conventional phenotype assignment, we included 561 subjects from the prospective CYPTAM study, which investigated the relationship between *CYP2D6* genotype and outcome of breast cancer treatment with tamoxifen (Supplementary data Fig.1)^23^. CYP2D6 enzyme activity can be inferred from the tamoxifen metabolism by using the ratio between the metabolites endoxifen and desmethyltamoxifen (Metabolic ratio (MR))^24^. To fully resolve the *CYP2D6* paternal and maternal alleles, we applied long-read sequencing^8^, yielding comparable predicted phenotype results to orthogonal testing (Supplementary data Fig.2 and Table 1). Classification of patients into conventional metabolizer categories resulted in 54.4% (R^2^=0.54) explained variability in CYP2D6 enzyme activity, (Fig. 1A, Supplementary data Table 2).

**Figure 1.**
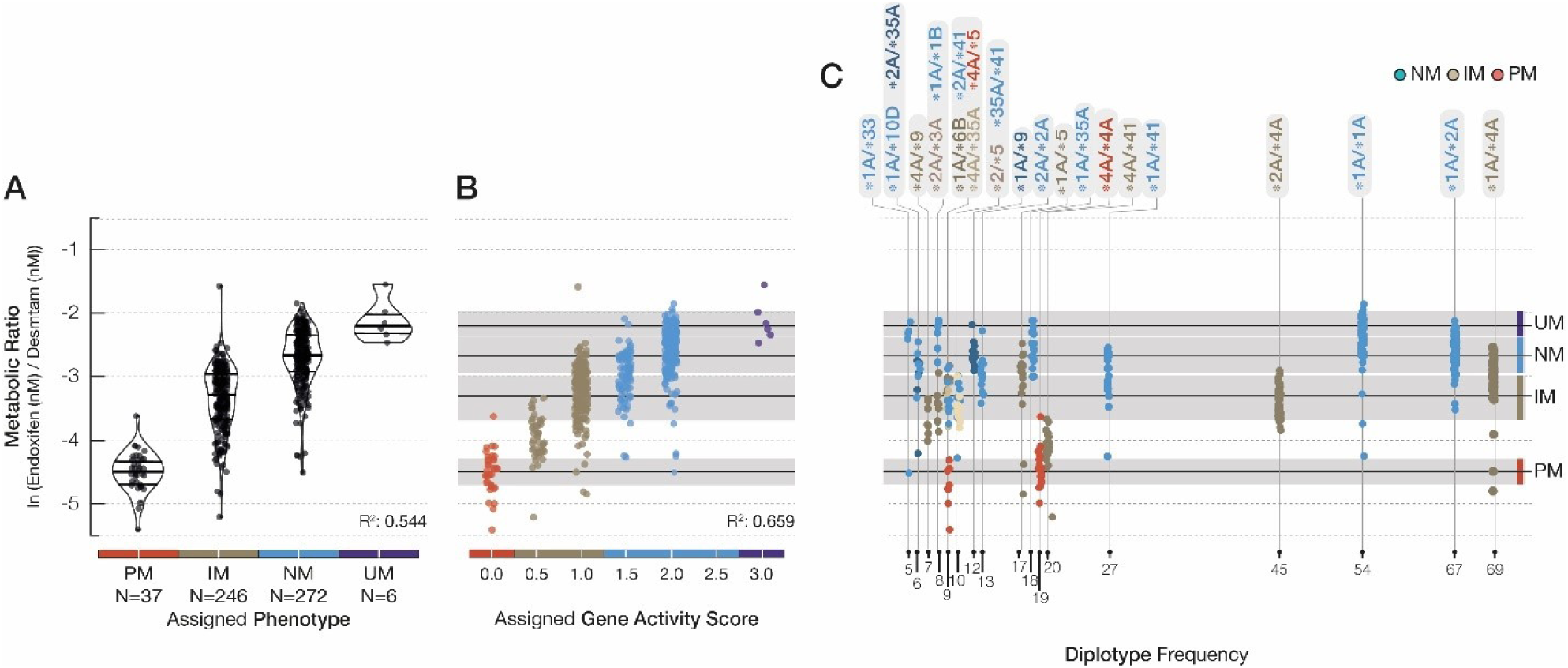
CYP2D6 activity based on conventional CYP2D6 metabolizer categories, gene-activity scores and diplotypes. Explained variability of CYP2D6 activity in the CYPTAM-cohort, based on (a) conventional phenotype categories and (b) gene activity scores (n=561). (c) range in enzyme activity within common (>1% occurrence) diplotypes. Metabolic ratio (ln(Endoxifen (nM)/Desmethyltamoxifen (nM)) serves as proxy for CYP2D6 enzyme activity. Gene activity scores and phenotype predictions are based on *-allele nomenclature and Dutch Pharmacogenetic Working Group translations using PacBio long-read sequencing data. R^2^: R^2^adjusted based on linear regression. Violin plots display observation density, lines represent the median and inter quartile range in (a). in (b and c): black lines represent median, grey area represents 95%Confidence interval. PM: Poor Metabolizer, IM: Intermediate Metabolizer, NM: Normal Metabolizer, UM: Ultra-rapid metabolizer

While the GAS system performs better than the 4 metabolizer categories (R^2^= 0.66), still a considerable amount of variability in enzyme activity within each predicted phenotype category remains unexplained (Fig.1B, Supplementary data Table 2). Stratifying the phenotype categories into diplotypes shows that the CYP2D6 activity varied substantially within identical diplotypes (Fig.1C). This suggests that a large proportion of the variability in enzyme activity within metabolizer phenotypes is already introduced when assigning haplotypes, with individuals carrying the same diplotype displaying phenotypes ranging from normal metabolizers to poor metabolizers.

## A continuous scale improves phenotype predictions

In order to increase the explained variability in CYP2D6 enzyme activity, we developed and trained a neural network consisting of two parts (Supplementary data Fig.3). The first level assigns contribution scores to individual alleles and variants while the second level combines paternal and maternal allelic scores into a predicted MR. Both parts were trained simultaneously on data generated from the CYPTAM cohort. By including all observed variants independent of predefined haplotypes, the explained variability increased to 79% (R^2^-adjusted = 0.79 (Fig.2A, Supplementary Table 2)). Inter individual variability is reflected by the range of observed MR in individuals with the same genetic make-up (equal predicted MR).The error rate (|observed MR – predicted MR|) was consistent over the range of the measured phenotype, with the exception of several (16 (2.9%) outside of confidence interval) subjects with a lower observed CYP2D6 activity than predicted (Fig.2A).

**Figure 2:**
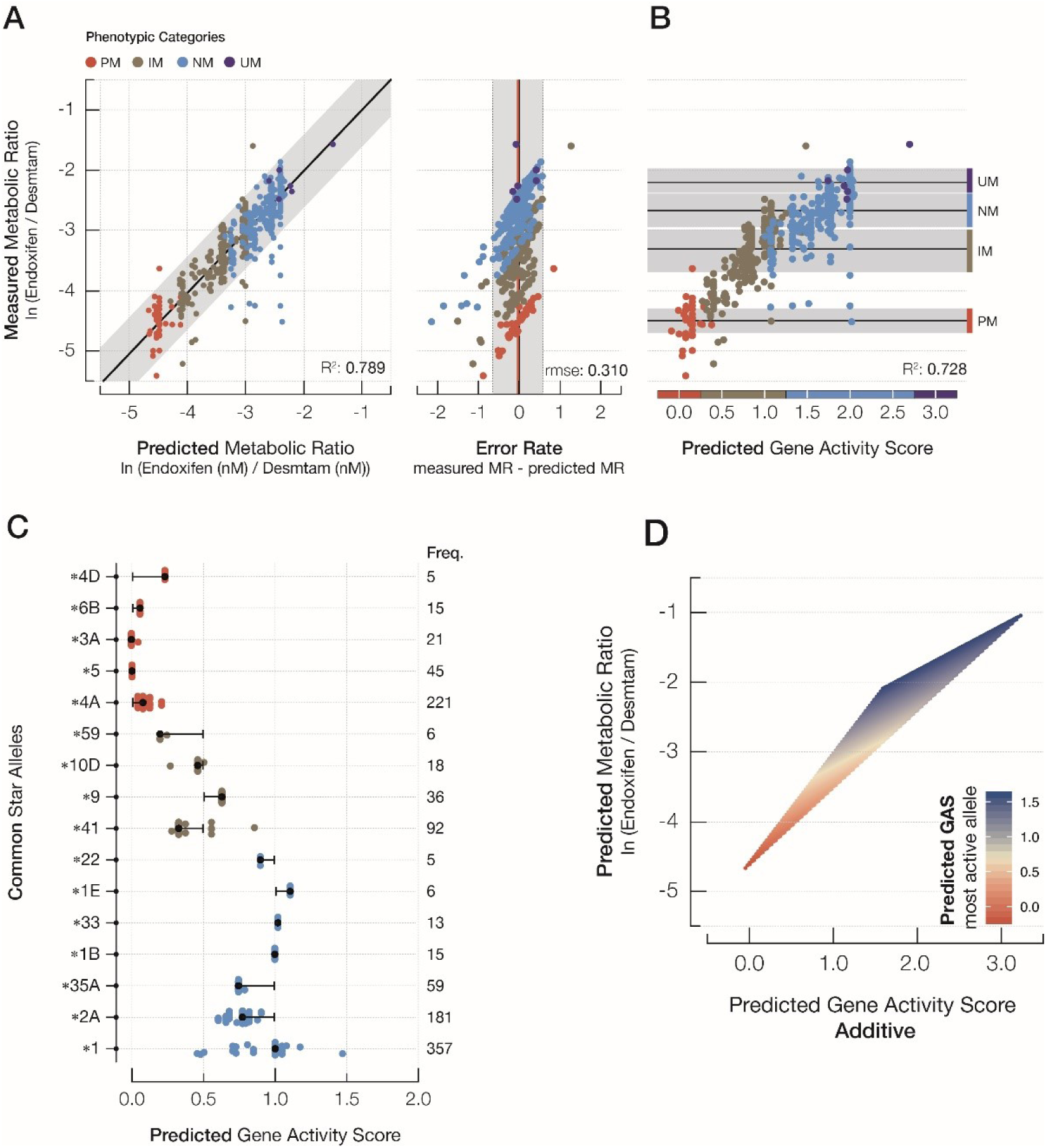
Neural network predictions for the CYPTAM-cohort. (a) The model predicts the metabolic ratio (ln(Endoxifen(nM)/Desmethyltamoxifen(nM)) as a proxy for CYP2D6 enzyme activity on a continuous scale, with a consistent error rate over the entire range (N=561). (b) the explained variability in enzyme activity using an additive model for the predicted allele contributions (N=561), e.g. predicted gene activity score = predicted contribution allele 1 + predicted contribution allele 2. (c) Predicted contributions per allele grouped in conventional *-allele assignments, (d) comparison of the neural network predicted gene activity score in an additive model with the neural network predicted metabolic ratio. Where the predicted gene activity score additive = predicted contribution allele 1 + predicted contribution allele 2, the predicted metabolic ratio is the final outcome of the neural network and the colours represent the activity of the most active allele. The neural network output suggest that the predicted metabolic ratio is not represented by the predicted gene activity scores in an additive model, but is better described by a neural network based non-additive relation. R^2^: R^2^adjusted based on linear regression. Rmse: root mean square deviation. In (a) and (b): blacklines represent median, grey area represents 95% Confidence interval MR: Metabolic Ratio (of ln(Endoxifen(nM)/ desmethyltamoxifen(nM))), PM: poor metabolizer, IM: Intermediate Metabolizer, NM: Normal Metabolizer, UM: Ultra-rapid Metabolizer

Allele contribution scores generated in the first part of the model were scaled to be comparable to the conventional GAS system (ranging 0-2). Interestingly, allele contribution scores predicted by the model showed a deviation from the conventional GAS assignments for multiple *-alleles (Fig.2C).

For example, the *2A allele has a conventional GAS of 1.0 representing a fully active allele. However, the predicted allele contributions ranged from 0.60 to 0.90, accounting for variants which are not included in the reference *2A haplotype. Similarly, the predicted average contribution for *41 is 0.34 (95% CI: 0.33-0.36), while the conventional assignment for the *41 allele is 0.5 ^12^. The same holds for the relatively rare *59 allele, currently regarded as decreased activity assuming a GAS of 0.5 whereas the activity is predicted to be 0.20 (95%CI: 0.19-0.22). The use of allele contribution scores on a continuous scale in an additive model improves the prediction of enzyme activity to 73% (Fig.2B). However, simply applying an additive model to individual allele contribution scores may be an oversimplification of human physiology. The second part of the neural network can accommodate non-additive combinations and therefore identify more complex relations between 2 alleles. Indeed, when the sum of allele contributions remains the same, a higher overall activity was observed when one of the alleles was fully active and one was fully inactive compared to two alleles with decreased activity (Fig.2D), which is in concordance with previous reports on IM phenotype variability^25^. These results can either be explained by a genetic component (e.g. up regulation, non-CYP2D6 variants) or by a non-linear relation between CYP2D6 enzyme activity and the metabolic ratio. There is, however, no indication to assume a non-linear relation in CYP2D6 enzyme activity and tamoxifen metabolism^26,27^.

## Increased explained variability in independent samples and CYP2D6 substrates

To validate the model, 167 subjects receiving tamoxifen who participated in the CYPTAM-BRUT study^28^, were sequenced using long reads and analysed with our neural network (Supplementary data Fig.1 and Table 1). In this cohort, patients are divided in two groups based on the use of CYP2D6 inhibitors that could influence the measured metabolic ratio of tamoxifen. Conventional phenotype predictions explained only 34.9% (R^2^-adjusted = 0.35 of the variability in CYP2D6 enzyme activity (Fig.3A and Supplementary data Table 2) in subjects without concomitant CYP2D6 inhibiting drugs (N=127). The deviation from the CYPTAM cohort can be explained by the significantly lower sample size. Neural network-based phenotype prediction on a continuous scale resulted in an almost doubling of the explained variability (R^2^-adjusted = 0.66) (Fig.3B and Supplementary data Table 2). For subjects using concomitant CYP2D6 inhibitors (N=24), 7.7% and 16.3% could be explained by conventional and continuous phenotype prediction respectively (Supplementary data Table 2). Moreover, there is a substantial overlap between subjects with a negative deviation from the predicted enzyme activity in the CYPTAM cohort and patients from the CYPTAM-BRUT cohort receiving concomitant treatment with a CYP2D6 inhibitor. This observation suggests that the concomitant use of CYP2D6 inhibitors may explain the overestimated enzyme activity for several subjects in the CYPTAM cohort. Stratifying the error between the predicted and observed enzyme activity per CYP2D6 inhibiting drug provides an estimate of the potency of the inhibitor (Fig.3B). In our data, paroxetine is identified as most potent CYP2D6 inhibitor, a finding which is supported by literature^29^.

**Figure 3.**
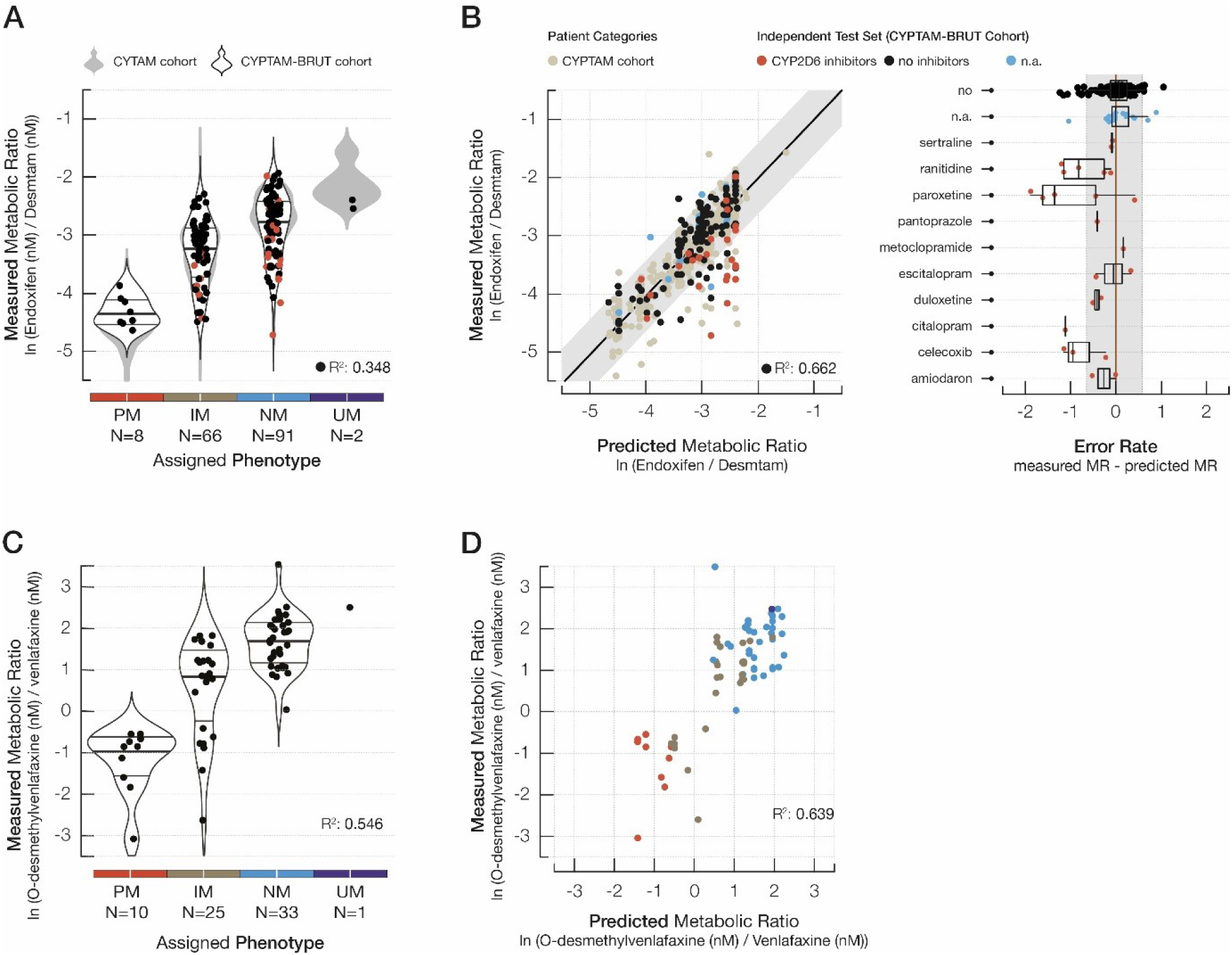
Conventional and continuous predictions in replication cohorts. The explained variability of CYP2D6 enzyme activity based on conventional phenotype categories for (a) CYPTAM-BRUT (Tamoxifen metabolism, N=167) and the (c) Venlafaxine-cohort (venlafaxine metabolism, N=69) and based on a neural network trained with data from the CYPTAM-cohort (b, d). (b) The influence of CYP2D6 inhibiting drugs on the overall enzyme activity shows overlap with CYPTAM samples with an negative deviation from the predicted enzyme activity. Error rates per CYP2D6 inhibiting drug give an indication of inhibitor potency (b, N=24 total). R^2^: R^2^adjusted based on linear regression. In (b) and (d): blacklines represent median, grey area represents 95% Confidence interval Violin plots display observation density, lines represent the median and inter quartile range. PM: poor metabolizer, IM: Intermediate Metabolizer, NM: Normal Metabolizer, UM: Ultra-rapid Metabolizer.

CYP2D6 enzyme activity is substrate specific and the effect on metabolism of a given variant varies per drug^30,31^. To assess substrate specificity of the neural network, we tested the performance on patients treated with a different CYP2D6 specific substrate, the antidepressant venlafaxine^32^ (Supplementary data Fig 1, Table 1 and Table 2). In venlafaxine treated patients, the explained variability of CYP2D6 activity increased from 54.6% (R^2^-adjusted = 0.55) for conventional phenotype prediction to 63.9% (R^2^-adjusted = 0.64) for phenotype predictions on a continuous scale (Fig.3C-3D). While the explained variability improves with the continuous prediction compared to conventional categorization, the increase is limited. Both substrate specificity^30,31^ as well as the limited sample size (N=69) in the venlafaxine can contribute to this limited increase in explained variability. Due to the limited sample size the majority of samples are from the intermediate and normal metabolizer phenotype, limiting the genetic diversity and thereby the R^2^.

## In vitro validation of predicted variant contributions

Furthermore, the trained neural network is queried to assess the contribution of individual variants to the overall enzyme activity (Supplementary data Table 3). These contributions show a wide range of effects in a pattern indicative of a continuous scale as opposed to an on/off effect (Fig.4A). To confirm the contributions predicted by the model, 4 variants and the *2 allele are expressed in HEK293 cells and incubated with bufuralol (Supplementary data Table 4). The direction of the predictions (decrease or increase) match the in vitro results, with an increased accuracy as the variant frequency increases. The predicted activity for *2 perfectly matched the observed activity in HEKcells (Fig.4B). Interestingly, for *CYP2D6*\*2 it is known that the normal activity of the allele is generally caused by the presence of an enhancer mutation causing an increase in expression which is almost in full linkage disequilibrium with the *2A allele^33,34^. In in vitro testing this enhancer mutation is not included resulting in a lower activity compared to wildtype. The presence of the enhancer mutation was not included in the neural network model, but was present in the population^35^. Nonetheless, in this study we observed a decreased *CYP2D6*\*2A activity in vivo, suggesting that *2A activity might be substrate dependent.

**Figure 4:**
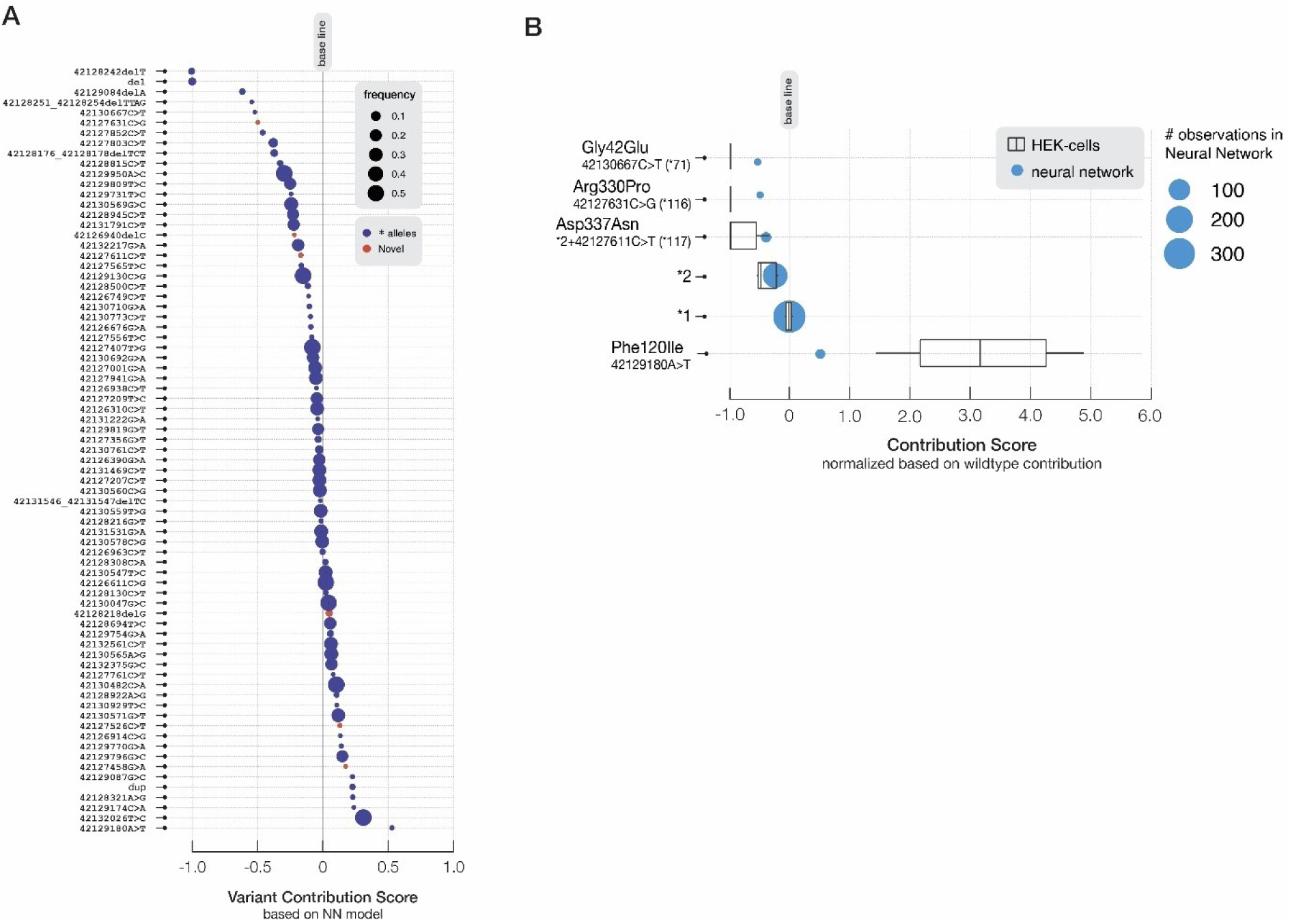
Contributions of individual variants. (a) Predicted contributions per variant included in the training of the Neural Network, with the absence of the variant set to 0.0 and a gene deletion set to −1.0 (n=78 variants). (b) in vitro validation of variants in HEK cells using bufuralol metabolism. Rate of bufuralol metabolism was normalized using the metabolic rate of the cells transfected with CYP2D6*1 cDNA as reference point of 1.0, similar to the neural network contributions results were further scaled to have the absence of a variant (normal activity a) set to 0.0 and the full absence of metabolism to −1.0. Incubations were performed in quadruple.

For the gain of function variant Phe120Ile, the difference was 8-fold, which might be explained by both the low frequency in the training cohort (n=3) and the substrate specificity of this variant^31,36,37^. The Phe120Ile amino substitution has previously been seen in *CYP2D6*49* where the allele has lower activity due to the presence of the Pro34Ser mutation^12^ and in *CYP2D6*53* where the Phe120Ile mutation is accompanied by an Ala122Ser. These alleles were not present in our study population, as none of the subjects carried the other mutations in combination with the Phe120Ile mutation^31^. These results indicate that the model was able to detect the biological effect of variants with improved accuracy as the number of observations increase (Fig.4B).

Here we have shown that the use of a continuous phenotype prediction based on a neural network and full gene sequencing data significantly decreases the missing heritability in CYP2D6 mediated metabolism. Our neural network is agnostic of the conventional GAS and *-alleles and does not assume an additive model for allele contributions to CYP2D6 enzyme activity as is the current standard. While the model shows generalizability to other CYP2D6 substrates, the venlafaxine data as well as the in vitro data also indicate the importance of substrate specificity, potentially requiring the development of gene-drug specific neural network prediction models.

More accurate predictions may improve the clinical impact of currently applied pharmacogenomic tests and help to optimize treatment of individual patients. A similar approach to what was used in this study could be implemented for improved phenotype prediction for other drug metabolizing enzymes of the CYP450 family. The impact of concomitant use of CYP2D6 inhibitors is highlighted by the low explained variability in patients using these drugs, reflecting the model’s ability to identify biological processes by identifying external factors as outliers and emphasizing the models capability of reflecting only the genetic impact on CYP2D6 activity.

After 30 years of *-allele and category-based pharmacogenomics, our continuous phenotyping approach paves the way for a new era of personalized medicine using advanced sequencing technologies and machine learning methods to improve prediction of variable drug response.

## Methods

### Study cohorts

The data used in this study originated from one main cohort and two independent cohorts. The main study cohort, the CYPTAM-cohort, consisted of 608 subjects for whom DNA material was available (The Netherlands National Trial Register: NTR1509)^1^. In short, the multicenter prospective CYPTAM study recruited subjects receiving tamoxifen as an adjuvant breast cancer therapy to investigate the association between *CYP2D6* genotype, endoxifen serum concentration and clinical outcomes. The first replication cohort, the CYPTAM-BRUT cohort, consisted of 225 subjects recruited in a study investigating the association between *CYP2D6* genotype and endoxifen serum concentration on response rate to tamoxifen in postmenopausal women (Clinicaltrials.gov: NCT00965939)^2^. The second replication cohort, the Venlafaxine-cohort, consisted of 78 subjects using venlafaxine. Samples were collected as part of routine patient care at the Catharina Hospital, Eindhoven, the Netherlands. DNA samples and accompanying data were de-identified before transfer to the Leiden University Medical Center (LUMC) for analysis.

### Drug metabolite measurements

For both the CYPTAM and CYPTAM-BRUT cohort, steady state through levels of tamoxifen and metabolites were measured with a validated high performance liquid chromatography-tandem mass spectrometry upon study inclusion. All measurements were performed at the LUMC department of Clinical Pharmacy and Toxicology. In total 4 compounds were measured: tamoxifen, 4-hydroxytamoxifen, O-desmethyltamoxifen and endoxifen. A total of 0.2ml of each serum sample was mixed with 0.5ml of 0.1M ZnSO4 and 0.2ml of the internal standard working solution 4-D5-IS. After mixing for 3min on a vortex mixer, the mix was centrifuged at 13000rpm for 5min at room temperature. A volume of 20μl supernatant was injected into the HPLC instrument. Chromatographic analysis was performed using a Waters Micromass Quattro micro API Tandem MS equipped with a Dionex P680A DGP-6HPLC pump, Dionex Ultimate 3000 autosampler and a Diones Thermostated Column Compartment. Separation of the analytes from potentially interfering serum components was achieved using a Waters X-bridge Column (3.5μm, 4.6 × 50mm) with a Spark HySphere C18 HD pre-column (7μm) in a Phenomenex holder. The mobile phase consisted of 25% solution A (0.1% formic acid + 2mM ammonium acetate in H2O) and 75% of solution B (0.1% formic acid + 2mM ammonium acetate in methanol) and was delivered at a flow rate of 0.4ml/min. Concentrations were normalized to nM and metabolic ratios calculated to reflect the rate of conversion from one metabolite to the next. For the Venlafaxine-cohort, plasma concentrations of venlafaxine and its metabolite O-desmethylvenlafaxine were determined as part of routine clinical care. Concentrations were determined with a validated ultra-performance liquid chromatography-tandem mass spectrometry method. Clozapine-D4 dissolved in acetonitrile was used as internal standard in a concentration of 0.1 mg/L. To 100μl of each plasma sample, a volume of 300μl of internal standard solution was added and vortex-mixed for 30 seconds. After centrifugation for 10min at 10900rpm, a volume of 200ul of the supernatant was mixed with 200μl of a 5mM ammoniumacetate solution and 10μl of this mix was injected on the UPLC-MS/MS. Chromatographic analysis was performed using a Waters Acquity UPLC with a BEH C18 (2.1 x 100mm, 1.7μm) column at 40°C. The mobile phased consisted of 90% solution A (5mM ammoniumacetate + 0.05%formic acid) and 10% solution B (acetonitril 100%) and was delivered at a flowrate of 0.35ml/min. Concentrations were normalized to nM. All samples were analyzed at the Catharina hospital department of Clinical Pharmacy.

### DNA sample processing

DNA isolation was performed previously for the main studies and for routine clinical use. Remaining DNA samples were collected and transferred to the LUMC for sequencing. All samples were sequenced with Pacific Bioscience’s (PacBio) SMRT-sequencing technique using full length *CYP2D6* amplicons^3^. PacBio sequencing enables the identification of all variants in the locus, including those in difficult and repetitive regions in addition to obtaining fully phased paternal and maternal alleles^4^. To obtain *CYP2D6* amplicons, three separate two-step PCR reactions were executed, one for full length amplicons and two for Copy Number Variants (CNV) using a similar protocol to Buermans et al.^4^. All primers used were based on previous research by Gaedigk et al.^5,6^ and ordered from Integrated DNA Technologies (IDT)^7^ (Supplementary data Table 5).

The *CYP2D6* specific primers were designed to generate a 6.6.kB fragment covering the entire *CYP2D6* locus including upstream and downstream regions^5,6^. Target regions were amplified using the Takara LA Taq DNA polymerase kit (catalogus number RR002A)^8^. A 10μl reaction volume contained 50-100ng DNA, 1x PCR buffer with MgCl2, 0.4mM dNTPs, 0.4 μM of both of the full length *CYP2D6* primers and 0.4U Takara La taq. PCR cycle parameters were 3min at 95°C, followed by 30 cycles of 10sec 98°C and 15min 68°C, finished with 15min at 68°C. Subsequently, amplicon barcoding was performed using M13-tailed primers. These barcode primers were introduced in a second PCR with identical conditions to the first, using 1ul of the first PCR product and 5 cycles of amplification.

*CYP2D6* gene deletions were identified with a duplex PCR. The primer set consisted of *CYP2D6*-deletion specific primers and an internal control (IC)^5,6^. Target regions were amplified using the KAPA long range hotstart kit from kapa biosystems (REF: KK3502)^9^. The 10 μl reaction volume contained 50-100ng DNA, 0.5x PCR buffer, 1.7mM MgCl2, 0.3mM dNTPs, 0.5uM of *CYP2D6*-deletion specific primers, 0.375 μM of IC primers and 0.025U Kapa Hotstart polymerase. Cycle parameters were 3min at 95°C, followed by 30 cycles of 15sec 95°C and 10min 68°C.

*CYP2D6* gene duplication and *CYP2D6/CYP2D7* fusion gene conformations were identified using a triplex PCR protocol. The primer set contained the *CYP2D6* full length primers, *CYP2D6* duplication primers and *CYP2D6/CYP2D7* fusion gene primers. The 10 μl reaction volume contained 50-100ng DNA, 0.5x PCR buffer, 1.7mM MgCl2, 0.3mM dNTPs, 0.5μl DMSO, 0.5μl of the CYP2D6 full length forward primer and 0.75μl of the reverse primer, 0.375μl of both *CYP2D6-*duplication specific primers, 0.5μl of the CYP2D6 fusion gene primer and 0.025U Kapa Hotstart polymerase. PCR conditions were identical to the duplex PCR.

Presence of CNVs and fusion genes was assessed on a 0.7% agarose gel with ethidium bromide staining, set at 100mV with a 55min run time. CNV and fusion gene positive samples, identified as additional fragments besides a full length or IC fragment, were selected for the subsequent barcoding PCR. For the selected samples of both the duplex and triplex PCR, barcoding was done with M13-tailed primers. Identical conditions to the first PCR were used with 1ul of PCR product from the first PCR and 5 cycles of amplification.

Barcoded amplicons were equimolar pooled into a full-length pool and a CNV and fusion genes pool. For CYPTAM, one pool of full-length samples per 96 wells plate was made and one pool for all CNVs and fusion genes. For CYPTAM-BRUT and the Venlafaxine-cohort, one pool with all full-length samples and one pool for all CNV and fusion gene samples of both cohorts was made. All pools were concentrated with Ampure XP beads (Agencourt). For the full-length fragment, additional size-selection was performed using BluePippin (Sage Science) to remove all fragments shorter than 5kB prior to pooling with the CNV and fusion gene amplicons. SMRTbell library preparation was performed on 500ng purified and size-selected PCR pool following the procedure & checklist – Amplicon template preparation and sequencing (PN 100-801-600 Version 04, Pacific Biosciences) and using SMRTbell template Prep Kit 1.0-SPv3^10^. The final SMRT library was sequenced on the PacBio RSII for the CYPTAM cohort and on the PacBio Sequel for the replication cohorts. For RSII, libraries were sequenced using sequencing primer V2 and P6-C4 chemistry with a movie time of 6hr, with a maximum of 96 samples per SMRT cell^10^. For Sequel, libraries were sequenced using sequencing primer V3, sequencing kit 3.0 and binding kit 3.0 on a 1M v3 LR SMRT cell with a movie time of 20hr, with a maximum of 288 samples per SMRT cell^10^. Deletions, duplications and hybrids were analyzed on a separate SMRT cell for all cohorts.

### Data preprocessing

The full pipeline for downstream processing is available at https://github.com/lumc-pgx/pgx-pipe. All downstream processing was run on a high performance computing cluster running the sun grid engine. Raw sequences were demultiplexed using LIMA followed by the CCS tool to generate CCS sequences. The subsequent haplotype phasing was performed using a custom pipeline which utilizes the CCS sequences to identify molecules originating from the same allele. Subreads of the CCS sequences were used to generate high quality phased allelic sequences per allele per individual using subreads of all molecules belonging to the same allele. Allelic sequences showing signs of disjoint sequences or chimeras were flagged. Per subject all phased allelic sequences were saved and plotted based on genomic distance.

Phased sequences were aligned to the *CYP2D6* sequence from GRCh38 and variants were called. A semi-global alignment was performed using biopython pairwise2, alignments were polished to ensure consistent indel positioning. Pharmacogenomic haplotype assignments were made based on PharmGKB translation tables^11^. For all haplotypes, the *-allele with a perfect match based on all variants observed was assigned, where the number of variants is decisive in the case of multiple perfect matches. When no perfect match is found the *1 haplotype was assigned. All identified variants were run through VEP (variant effect predictor) to determine their potential impact on protein function^12^. Variants were flagged as ‘known’ for variants in *-allele nomenclature, ‘novel’ for variants not in *-allele nomenclature, ‘in polymer region’ for variants located in homo-polymer regions.

The phased alleles were separated from chimeras and disjoint sequences by manual curation based on genomic distance plots and the presence of chimeras and disjoint flags. A cut-off of at least 10 molecules per allele and 10 passes per molecule was used to determine the reliability of the sequences. In the presence of gene deletion, the second allele was annotated as ‘deletion’. In the case of a duplication, the duplicated allele, determined based on the number of molecules observed per allele, was annotated as *‘*duplicated’. Subjects identified as carrying a *CYP2D6/2D7* fusion gene were annotated as ‘hybrid’ for the hybrid allele. Selected alleles were linked to the clinical data based on subject specific barcodes, resulting in one datafile per cohort containing clinical data, selected alleles and haplotype calls.

### Prediction models

For further analysis, samples were selected based on the presence of full length *CYP2D6* sequences, the absence of *CYP2D6/CYP2D7* conversions and fusion genes, and on the presences of clinical data regarding drug metabolism (N=561 for CYPTAM, N=167 for CYPTAM-BRUT, N=69 for Venlafaxine). For each cohort the clinical datasets containing metabolite levels were merged with the sequencing data containing the assigned haplotypes.

#### Conventional method

For the CYPTAM-cohort, haplotype and phenotype assignments based on PacBio sequencing data were compared to calls from the Roche Amplichip which were determined previously. To assess explained variability based on conventional phenotyping, the same methods were applied to all three cohorts. For the CYPTAM-BRUT cohort data on concomitant use of CYP2D6 inhibiting drugs was available, based on which the cohort was split into ‘non-inhibitor users’, ‘inhibitor users’ and ‘unknown inhibitor use’.

For all cohorts the same methods were applied. Haplotype calls were translated into Gene Activity Scores (GAS) and predicted phenotype categories based on the CPIC and DPWG guidelines^13,14^. A GAS of 0.0 was assigned to non-active alleles, 0.5 to decreased activity, 1.0 to normal activity and 2.0 to increased activity alleles. Subsequently the scores per allele were combined into the overall GAS, followed by a translation into phenotype categories. Based on the guidelines 4 clinically implemented phenotype categories were assigned: poor metabolizer (PM, GAS = 0.0), intermediate metabolizer (IM, GAS =0.5-1.0), normal metabolizers (NM, GAS=1.5-2.5) and ultra-rapid metabolizers (GAS = 3.0).

As a proxy for CYP2D6 enzyme activity, the metabolic ratio of the most CYP2D6 specific conversion of either tamoxifen or venlafaxine metabolism was used. For the CYPTAM and the CYPTAM-BRUT cohorts, the log of the metabolic ratio of the conversion from desmethyltamoxifen to endoxifen ((LN(Endoxifen(nM)/ Desmethyltamoxifen(nM))) was used as a proxy for CYP2D6 enzyme activity^14,15^. Transformation to log was performed to normalize the data. There are no indications to assume non-linear kinetics of endoxifen formation by CYP2D6^16^, in fact the kinetics of all other metabolites are shown to be perfectly linear^17^. Additionally, the metabolic ratio as used in this study was shown to stay consistent with dose increase for all phenotypes^18,19^ making it a suitable proxy for enzyme activity. Finally, it is expected that intra-individual variability of CYP2D6 enzyme activity is limited, making one measurement at steady state a suitable approach ^20-22^.

For the Venlafaxine cohort, the log of the metabolic ratio for the conversion from venlafaxine to desmethylvenlafaxine (LN(O-desmethylvenlafaxine(nM) /venlafaxine(nM))) was used.

The amount of explained variability in CYP2D6 enzyme activity based on conventional phenotype predictions was assessed using linear regression, assuming a linear relation between predicted phenotypes and observed metabolic ratio. Two different models were assessed, the first based on the clinically applied phenotype categories (PM, IM, NM, UM), the second based on overall GAS. Explained variability was expressed as R^2^-adjusted, using a p<0.05 cutoff for significance.

#### Neural network

From the selected alleles per individual, a dataset was generated indicating the presence (1) or absence (0) of every variant observed in the entire cohort (including deletions and duplications). From these variants a selection is made, variants were excluded if they adhere to the following rules: located in homopolymer regions or not in *-allele nomenclature and synonymous, intronic, located upstream or downstream. These were excluded to prevent confounding from irrelevant variants in the development of the neural network. Variants were included if they were part of the *-allele nomenclature or if they were additionally nonsynonymous, frameshifts or splice sites variants.

The neural network was build using Keras (https://github.com/keras-team/keras) with the TensorFlow (https://github.com/tensorflow/tensorflow) backend. It uses the selected variants (extended data table 1) per allele as predictors (N=78) and the measured metabolic ratio (LN(Endoxifen(nM)/ Desmethyltamoxifen(nM))) as a surrogate for CYP2D6 enzyme activity and the outcome variable of the model. The model was comprised of 2 parts (Supplementary data Fig.3). The first consisted of two interpreters, one per allele, which train as one. These interpreters use all selected variants per allele as input data and combine them into an allele contribution. The second part was the combiner model which combined the two allele contributions to predict the metabolic ratio. The model was trained with the data from the CYPTAM cohort and trained both parts simultaneously. A 10-fold cross validation with 100 cycles both with and without internal hold-out was performed and showed no signs of overfitting (Supplementary data Fig.4). Shap (Shapley Additive explanation)-values were extracted and normalized to define allele contributions. Where 0.0 was assigned to a gene deletion and 1.0 to a fully wildtype allele. Variant contributions were normalized accordingly, resulting in the sum of variant contributions per allele corresponding to the allele contribution.

For both replication cohorts, the same variants as which were used during the training were included in the selection. For the Venlafaxine cohort, the predicted metabolic ratio is translated with a linear transformation into the metabolic ratio for venlafaxine (LN(O-desmethylvenlafaxine(nM)/venlafaxine(nM))). The explained variability for all cohorts was assessed using linear regression with the predicted metabolic ratio as predictor and the observed metabolic ratio as outcome. Explained variability was expressed as R^2^-adjusted, using a p<0.05 cutoff for significance. Error rate of the model was expressed as the rmse (root-mean-square error).

### In vitro validation

To confirm the contribution of individual variants as predicted by the neural network, four high impact variants and the *2 allele were tested in vitro. Variants were selected based on the following criteria: the predicted contribution had to be ≥0.2 or ≤ −0.2, there is no linkage disequilibrium with a known causal variant, the variant needs to be potentially causal (e.g. missense, frameshift), both gain of function and loss of function variants should be included. Variants selected were: g.42130667C>T, g.4212761C>G, g.42127611C>T and g.42129180A>T as well as the *2 allele. Site mutagenesis was performed on pCMV4 *CYP2D6*1* plasmid^23^ with QuikChange II Site-Directed Mutagenesis Kit (Agilent, CA, US). Plasmid cDNA encoding variants with the following amino acid exchanges were created: Arg330Pro (g.42127631C>G), Gly42Glu (g.42130667C>T) and Phe120Ile (g.42129180A>T). The Asp337Asn exchange was performed using pCMV4 *CYP2D6*2* as template. Mutagenesis primers and selected variants are listed in Supplementary data table 6. Variants were expressed in HEK293 cells, which were grown in DMEM 6046 (Sigma), containing 1g glucose/l, 10% fetal bovine serum and penicillin/streptomycin (100IU/ml, 100mg/ml) to a confluence of 60-70%. The pCMV4 vectors containing the variants were transfected using Viromer Red (Lipokalyx, Halle, Germany) according to manufacturer’s protocol. Cells were harvested after 24-48 hours incubation were stored at −80°C. Cell pellets were resuspended in 0.1M sodium phosphate buffer, followed by sonication for 20 x 1sec and were centrifuged at 800 x g. Incubations were performed with 800 x g supernatant corresponding to 25-125μg of protein, 0.1 M sodium phosphate buffer, 50 μM bufuralol (racemate) and 1 mM NADPH in a total volume of 150 μl. reactions were linear for at least 5 hours and were terminated by addition of 14μl of 70% perchloric acid. After centrifuging the supernatant was analysed by high performance liquid chromatography as described by Kronbach et al^24^. The levels of CYP2D6 apoprotein of the different allelic variants were determined using sodium dodecyl sulfate polyacrylamide gel electrophoresis and Western blot analysis. Residual CYP2D6 activity was assessed and normalized with the average activity of the *-allele set at 1.0 to allow for comparison with the neural network predictions.

### Software

All statistics were performed using R version 1.0.143, the haplotyping pipeline and neural network were developed using Python.

### Ethics

The CYPTAM protocol was approved by the Institutional Review board of the LUMC. The CYPTAM-BRUT protocol was approved by the Institutional Review board of the Leuven University medical center. Venlafaxine samples were collected in routine clinical care at Catharina Hospital, Eindhoven, the Netherlands and fully anonymized prior to analysis.

## Data availability statement

Sequence data (CYPTAM cohort) that support the findings of this study is in the process of being deposited in European Genome and Phenotype Atlas.

## Code availability statement

The code generated and used within this study is partially available on GitHub (https://github.com/lumc-pgx/pgx-pipe). Further code generated is not publicly available due to them containing information that could compromise research participant privacy/consent.

## Author contributions

Conception: SYA, HJG, JS

Design: ML, WGA, HJG, JS, SYA

Acquisition, analysis: ML, WGA, RM, RV, RB, BD, MD, PN, MIS, IJ, SG, SYA

Interpretation: ML, WGA, HJG, JS, SYA

Manuscript writing: ML, JS, SYA

All authors have approved the submitted version of the manuscript

