## Supplementary Fig. 1-4, tables 1,2,4-6 for "A unifying model to predict variable drug response for personalised medicine"

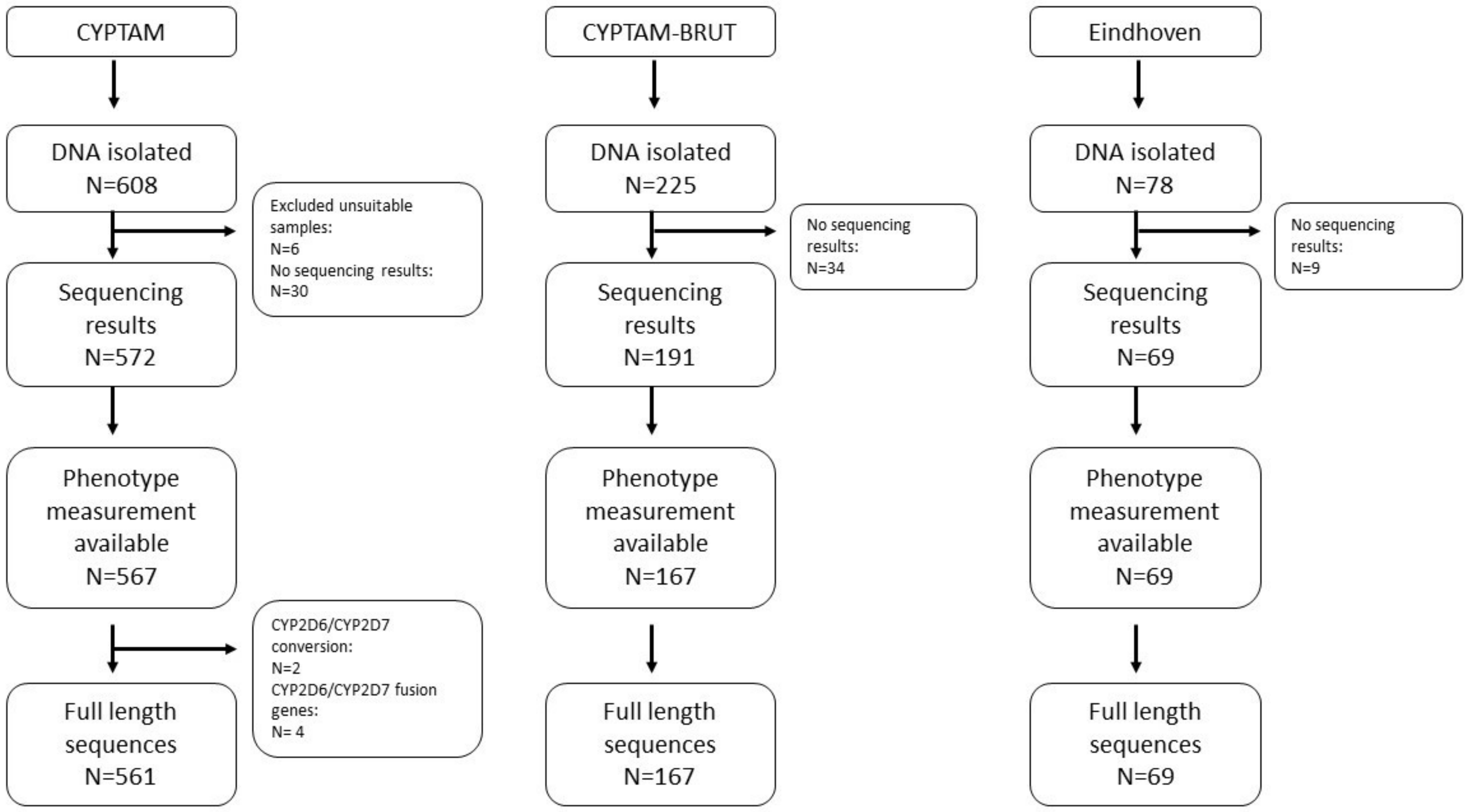

**Supplementary Figure 1 | Flowchart of Study cohorts.** Samples were selected based on availability of remaining DNA. Samples were excluded if patients no longer wanted to be part of the main study, or were double included (total: N=6 in CYPTAM). All samples were sequenced for *CYP2D6* with PacBio SMRT sequencing. For neural network training and predictions only samples with full length CYP2D6 sequences available and with clinical phenotype data were included.

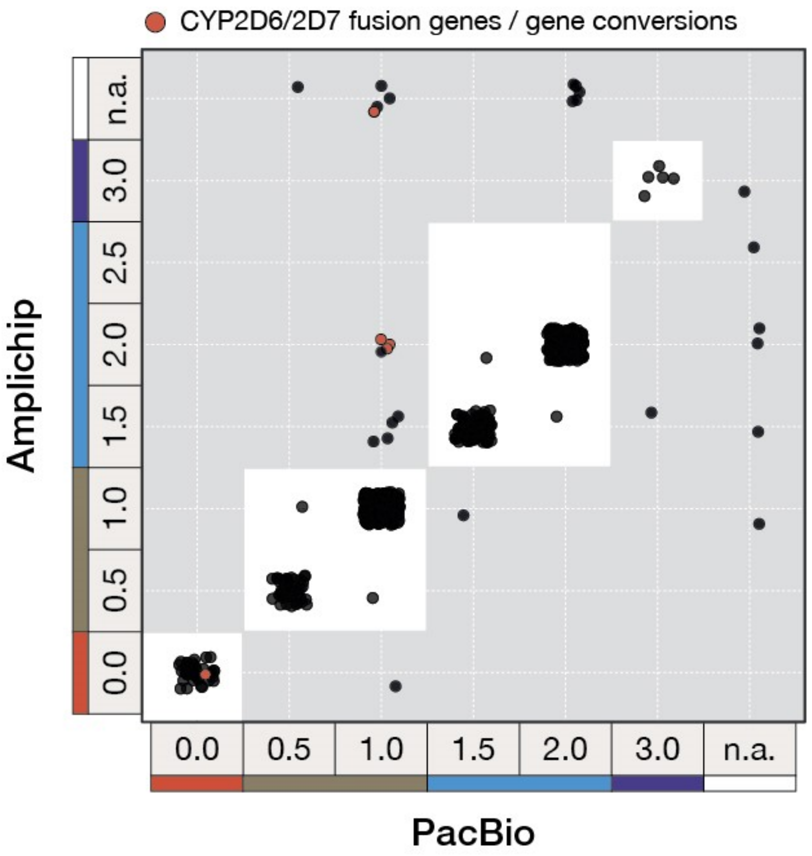

**Supplementary Figure 2 | Concordance Amplichip and PacBio based phenotype predictions.** For all CYPTAM individuals phenotype predictions were made, based on Amplichip genotyping and PacBio SMRT-sequencing genotype calling, according to Gene Activity Scores (GAS) and the Dutch Pharmacogenetics Working Group guidelines . Concordance was high: Kappa-coefficient: 0.94, p<0.0001

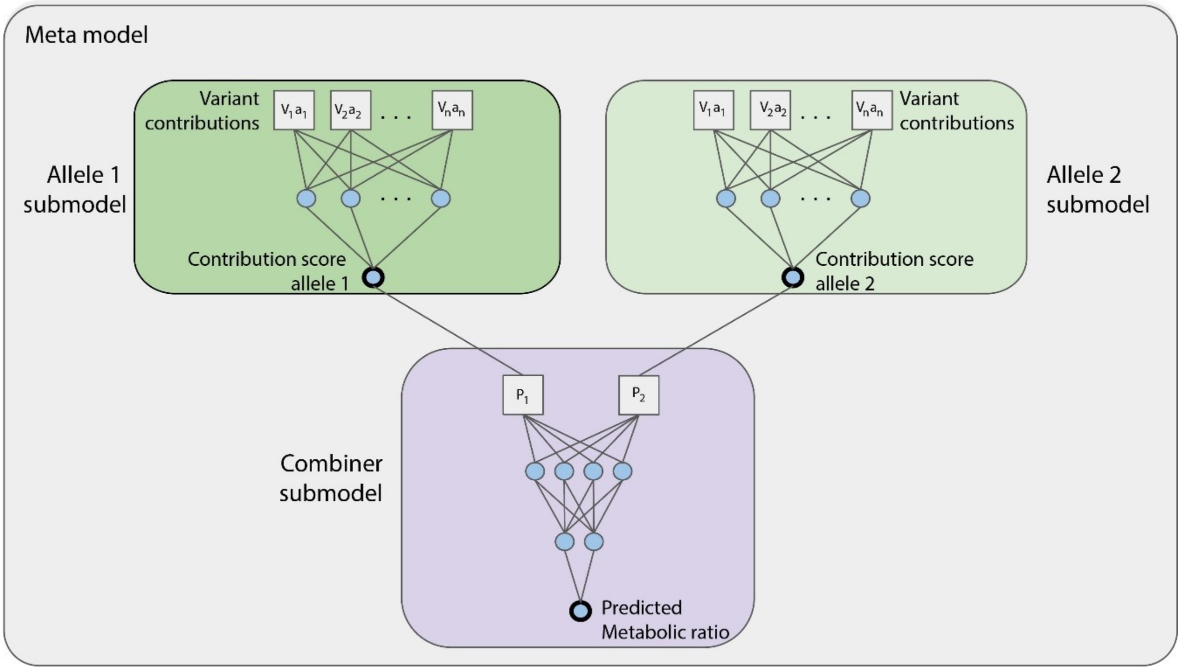

**Supplementary Figure 3 | Neural Network design.** The neural network model consists of two parts. First the allele submodels, one per allele, which train as one and result in contribution scores per allele. The second part, the combiner submodel, combines the contribution scores into a predicted Metabolic ratio. Variant contributions reflect the impact of variants on enzyme function, the contribution scores are normalized to represent gene activity scores, the predicted Metabolic ratio serves as a proxy for CYP2D6 enzyme activity.

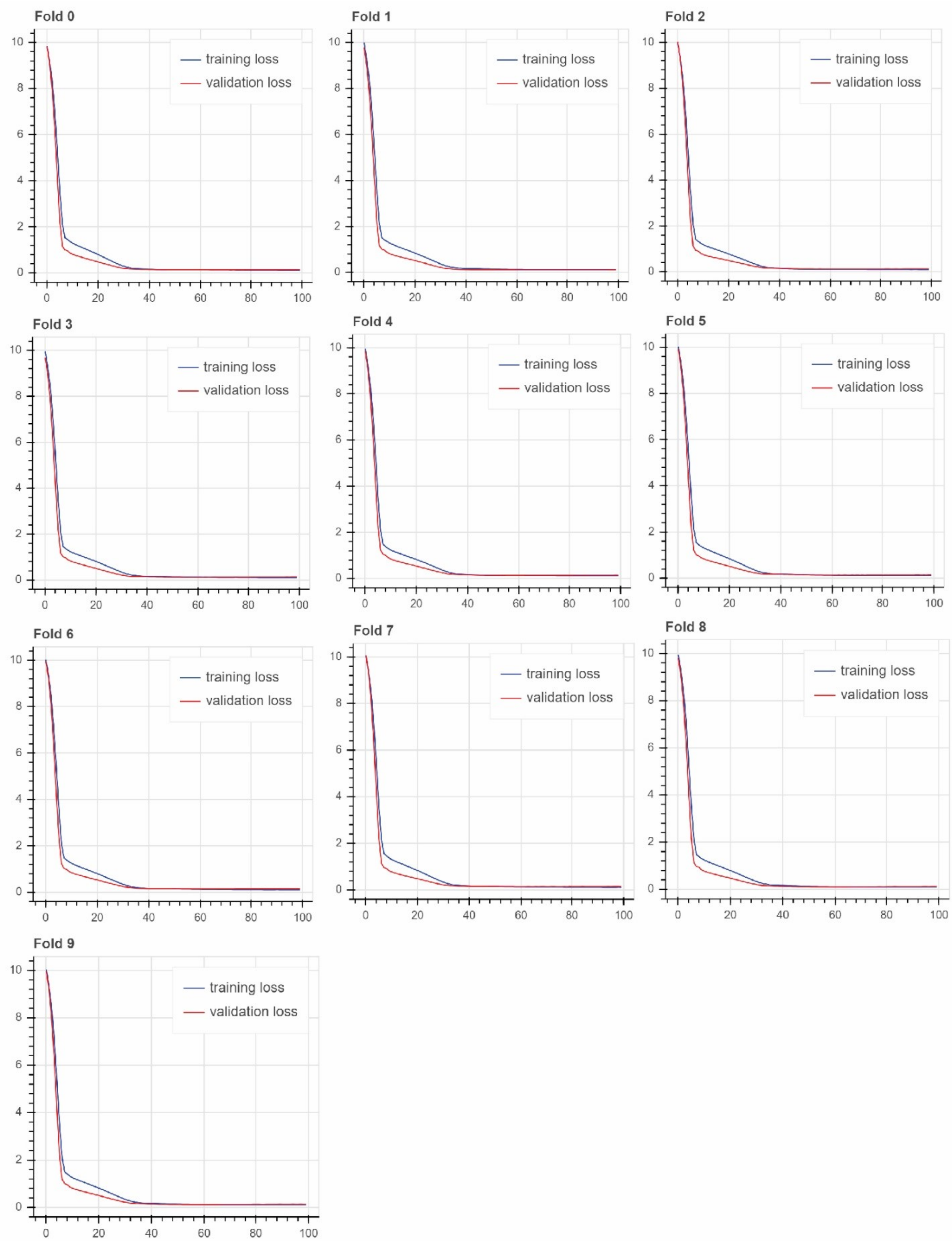

**Supplementary Figure 4 | 10-fold crossvalidtion with internal hold-out.** 25% of the data was used for the validation set. No devation between training and validation loss was observed  up to 100 epochs. No signs of overfitting were observed.

| **Haplotype** | **CYPTAM** | | **CYPTAM-BRUT** | | **Venlafaxine** | |
|---|---|---|---|---|---|---|
| | N | % | N | % | N | % |
| *1 | 357 | 32.8 | 78 | 30.7 | 33 | 23.91 |
| *108 | 3 | 0.27 | 1 | 0.39 | | |
| *10A | | | 1 | 0.39 | | |
| *10B | | | 1 | 0.39 | | |
| *10D | 18 | 1.6 | 3 | 1.2 | 2 | 1.45 |
| *15 | | | | | 1 | 0.72 |
| *17 | 2 | 0.18 | | | | |
| *1B | 15 | 1.3 | 2 | 0.79 | | |
| *1D | 2 | 0.18 | | | 1 | 0.72 |
| *1E | 6 | 0.53 | | | | |
| *1xN | 6 | 0.53 | 1 | 0.39 | | |
| *2 | 4 | 0.36 | | | | |
| *22 | 5 | 0.45 | | | | |
| *27 | | | 1 | 0.39 | | |
| *2A | 181 | 16.1 | 51 | 20.1 | 24 | 17.39 |
| *2AxN | 1 | 0.09 | 1 | 0.39 | 2 | 1.45 |
| *2D | | | 2 | 0.79 | 1 | 0.72 |
| *2M | 1 | 0.09 | | | | |
| *2xN | 1 | 0.09 | | | | |
| *31 | 1 | 0.09 | | | | |
| *33 | 13 | 1.2 | 1 | 0.39 | 2 | 1.45 |
| *34 | | | 1 | 0.39 | | |
| *35A | 59 | 5.3 | 18 | 7.1 | 8 | 5.80 |
| *35AxN | 2 | 0.18 | | | | |
| *39 | | | | | 1 | 0.72 |
| *3A | 21 | 1.9 | 3 | 1.2 | 7 | 5.07 |
| *41 | 92 | 8.2 | 24 | 9.4 | 16 | 11.59 |
| *41xN | 1 | 0.09 | | | | |
| *4A | 221 | 19.7 | 33 | 13.0 | 21 | 15.22 |
| *4AxN | 2 | 0.18 | 2 | 0.79 | 1 | 0.72 |
| *4B | | | | | 1 | 0.72 |
| *4D | 5 | 0.45 | 8 | 3.1 | 4 | 2.9 |
| *4H | 1 | 0.09 | | | | |
| *4J | | | 1 | 0.39 | | |
| *5 | 45 | 4.0 | 9 | 3.5 | 8 | 5.80 |
| *59 | 6 | 0.54 | | | | |
| *6A | | | | | | |
| *6B | 15 | 1.3 | 4 | 1.6 | 2 | 1.45 |
| *7 | | | 1 | 0.39 | | |
| *9 | 36 | 3.2 | 7 | 2.8 | 3 | 2.17 |

**Supplementary Table 1 | *-allele haplotype frequencies for all cohorts.** All haplotypes are based on all variants observed in the CYP2D6 locus, including CYP2D6/2D7 conversions and fusion genes. CYPTAM cohort is the training cohort, CYPTAM-BRUT the first validation cohort with tamoxifen as the CYP2D6 substrate used, Venlafaxine is the second replication cohort with individuals using the CYP2D6 substrate venlafaxine. Haplotype translations are based on the PharmGKB variant to haplotype translations.

| | **CYPTAM (N=561)** | **CYPTAM-BRUT no inhibitors (N=127)** | **CYPTAM-BRUT Inhibitors (N=24)** | **Venlafaxine (N=69)** |
|---|---|---|---|---|
| Categorical phenotype (DPWG) | 0.5443, P = 2.78 x $10^{-95}$ | 0.3483, P = 4.517 x $10^{-12}$ | 0.07752 P=0.1009 | 0.5461, P=8.09 x 10-$^{12}$ |
| Gene activity scores - conventional | 0.6589, P =3.63 x $10^{-128}$ | 0.4963, P=1.47 x $10^{-20}$ | 0.1583 P=0.0308 | 0.6326, P= 1.99 x $10^{-16}$ |
| Continuous phenotyping | **0.7885, P = 6.14 x $10^{-191}$** | **0.6618, P=1.99x$10^{-31}$** | 0.1633 P=0.0286 | **0.6385, P = 1.15 x $10^{-16}$** |
| Gene activity scores predicted - additive | 0.7278, P = 2.59 x $10^{-160}$ | 0.6012, P =6.15 x $10^{-27}$ | **0.1929, P=0.0183** | 0.6064, P = 2.028 x $10^{-15}$ |

**Supplementary Table 2 | Regression results explaining CYP2D6 enzyme activity using different methods.** All predicted phenotypes are based on PacBio long-read sequencing data. DPWG guidelines are used to assign the conventional four phenotype categories and gene activity scores. A neural network trained on CYPTAM data is used for the prediction of a continuous phenotype. Per allele contributions from the neural network are added together (similar to the gene activity score) to predict the effect of a continuous gene activity score per allele using an additive model. For CYPTAM-BRUT, patients with unknown inhibitor use were exclude from this analysis (N=16). DPWG: Dutch Pharmacogenetics Working Group.

| Genotype | September 2019 | | December 2019 | | Januari 2020 | | Average |
|---|---|---|---|---|---|---|---|
| *1 | 0,927845 | 1,072155 | 0,952799 | 1,047201 | 1,027143 | 0,972857 | 1 |
| *2 | 0,780427 | 0,806543 | 0,520376 | 0,474208 | 0,455362 | | 0,607383 |
| *2+D337N | 0,438914 | 0,66233 | 0 | 0 | 0,018975 | | 0,224044 |
| R330P | 0 | 0 | 0 | 0 | 0,007519 | 0,047079 | 0,0091 |
| G42E | 0 | 0 | 0 | 0 | 0,009874 | 0,001144 | 0,001836 |
| F120I | 3,439576 | 4,910881 | 5,878972 | 5,374999 | 3,100804 | 2,457602 | 4,193805 |

**Supplementary Table 4 | Bufuralol incubation results.** Site mutagenesis on HEK cells was performed on pCMV4 CYP2D6*1 plasmid with QuikChange II Site-Directed Mutagenesis Kit (Agilent, CA, US). The D337N exchange was performed using pCMV4 CYP2D6*2 as template. HEK cells were  were incubated with bufuralol for 5 hours, upon which the metabolism rate was assessed by measuring bufuralol and metabolites. Results are normalised to the average activity of *1, which is set at 1.0.

| Name | Primer Sequence (5’- 3’) |
|---|---|
| Full-length PCR | |
| Fragment A Forward | ATGGCAGCTGCCATACAATCCACCTG |
| Fragment A Reverse | CGACTGAGCCCTGGGAGGTAGGTAG |
| Duplex PCR (*5) | |
| Fragment *5-forward | CTCCAGCCTCCACCAGTCCAG |
| Fragment *5-reverse | CAGGCATGAGCTAAGGCACCCAGAC |
| IC-forward | GCATGCACAGCTCAGCACTGC |
| IC-reverse | GCCACCCTGATGTCTCAGTTTCG |
| Triplex PCR | |
| Fragment A Forward | ATGGCAGCTGCCATACAATCCACCTG |
| Fragment A Reverse | CGACTGAGCCCTGGGAGGTAGGTAG |
| Fragment B - Forward | CCATGGAAGCCCAGGACTGAGC |
| Fragment B – Reverse | CGGCAGTGGTCAGCTAATGAC |
| Fragment H - Forward | TCCGACCAGGCCTTTCTACCAC |

**Supplementary Table 5 | Primer sequences for three separate PCR reactions.** PCR Full length PCR: yielding one full length CYP2D6 sequence. PCR duplex (*5): yielding an internal control fragment for all samples and a deletion fragment if a CYP2D6 deletion is present. PCR triplex: Yielding a full length fragment as well as a duplication fragment in the presence of a CYP2D6 duplication and/or a hybrid fragment in the presence of a CYP2D6/CYP2D7 fusion gene. IC: Internal control

| Nucleotide mutation | Amino acid mutation | Primer forward 5´- 3 | Primer reverse 5´- 3 |
|---|---|---|---|
| 125 G>A | G42E | CCTGCCACTGCCCG**A**GCTGGGCAACCTGCT | AGCAGGTTGCCCAGC**T**CGGGCAGTGGCAGG |
| 1009 G>A | D337N | CAACAGGAGATCGAC**A**ACGTGATAGGGCAGG | CCTGCCCTATCACGT**T**GTCGATCTCCTGTTG |
| 1611 T>A | F120E | CGTTCCCAAGGGGTG**A**TCCTGGCGCGCTATG | CATAGCGCGCCAGGA**T**CACCCCTTGGGAACG |
| 3160 G>C | R330P | CGGATGTGCAGCGCC**C**TGTCCAACAGGAGAT | ATCTCCTGTTGGACA**G**GGCGCTGCACATCCG |

**Supplementary data table 6 | HEK cell mutagenisis primers.** Site mutagenesis on HEK cells was performed on pCMV4 CYP2D6*1 plasmid with QuikChange II Site-Directed Mutagenesis Kit (Agilent, CA, US). The D337N exchange was performed using pCMV4 CYP2D6*2 as template
