## Supplementary table 3 for "A unifying model to predict variable drug response for personalised medicine"

**Supplementary data table 3: Variant frequencies and predicted contributions.** All variants (mapped to GRCh38) which are identified in all individuals included in this study. Variant Effect Prediction (VEP) based on the most severe effect are included. Variants were included in the neural network if they were part of *-allele nomenclature or if they were classified as missense, frameshift or splice region variants. Based on the neural network each variant was assigned a variant contribution score scaled to -1.0 for a deletion and 0 for no effect compared to wildtype.

| Variant (NC_000022.11:g.) | CYPTAM | BRUT | EIND | Total | Minor allele frequency (1596 alleles) | included neural network | Predicted contribution to allele activity based on neural network | consequence | rs-number | sift prediction | sift score | polyphen prediction | polyphen score |
|---|---|---|---|---|---|---|---|---|---|---|---|---|---|
| 42126069A>C | 600 | 187 | 81 | 868 | 54,4% | no | | downstream_gene_variant | rs35028622 | - | - | - | - |
| 42126074G>A | 2 | 0 | 0 | 2 | 0,1% | no | | downstream_gene_variant | rs77827855 | - | - | - | - |
| 42126079C>T | 2 | 0 | 0 | 2 | 0,1% | no | | downstream_gene_variant | rs4078249 | - | - | - | - |
| 42126136_42126138delTGT | 348 | 126 | 51 | 525 | 32,9% | no | | downstream_gene_variant | rs71184866 | - | - | - | - |
| 42126136delT | 3 | 0 | 0 | 3 | 0,2% | no | | downstream_gene_variant | | - | - | - | - |
| 42126138_42126139delTG | 6 | 0 | 0 | 6 | 0,4% | no | | downstream_gene_variant | | - | - | - | - |
| 42126201_42126202insG | 0 | 1 | 0 | 1 | 0,1% | no | | downstream_gene_variant | | - | - | - | - |
| 42126310C>T | 348 | 127 | 52 | 527 | 33,0% | yes | -0.043653817811100566 | downstream_gene_variant | rs12169962 | - | - | - | - |
| 42126343_42126344insG | 1 | 0 | 0 | 1 | 0,1% | no | | downstream_gene_variant | rs1200940710 | - | - | - | - |
| 42126347T>C | 1 | 0 | 0 | 1 | 0,1% | no | | downstream_gene_variant | rs148648640 | - | - | - | - |
| 42126389_42126390insA | 0 | 2 | 0 | 2 | 0,1% | no | | downstream_gene_variant | | - | - | - | - |
| 42126390G>A | 247 | 60 | 29 | 336 | 21,1% | yes | -0.027091746508636685 | downstream_gene_variant | rs28371738 | - | - | - | - |
| 42126396_42126397delCT | 2 | 0 | 0 | 2 | 0,1% | no | | downstream_gene_variant | rs202032066 | - | - | - | - |
| 42126417G>A | 1 | 0 | 0 | 1 | 0,1% | no | | downstream_gene_variant | rs550569325 | - | - | - | - |
| 42126462G>A | 14 | 2 | 0 | 16 | 1,0% | no | | downstream_gene_variant | rs28371737 | - | - | - | - |
| 42126552G>T | 0 | 0 | 2 | 2 | 0,1% | no | | 3_prime_UTR_variant | rs746767060 | - | - | - | - |
| 42126611C>G | 601 | 184 | 81 | 866 | 54,3% | yes | 0.02252113634360636 | missense_variant | rs1135840 | tolerated | 1 | benign | 0 |
| 42126611delC | 0 | 1 | 0 | 1 | 0,1% | no | | frameshift_variant | | - | - | - | - |
| 42126676G>A | 5 | 2 | 0 | 7 | 0,4% | yes | -0.09213514232176667 | synonymous_variant | rs150445731 | - | - | - | - |
| 42126749C>T | 1 | 0 | 0 | 1 | 0,1% | yes | -0.109389898471713 | missense_variant | rs267608319 | deleterious | 0.04 | benign | 0.255 |
| 42126763G>T | 2 | 0 | 0 | 2 | 0,1% | no | | intron_variant | rs548264542 | - | - | - | - |
| 42126798A>G | 0 | 1 | 0 | 1 | 0,1% | no | | intron_variant | rs1243469456 | - | - | - | - |
| 42126914C>G | 1 | 0 | 0 | 1 | 0,1% | yes | 0.133014501516827 | missense_variant | rs28371733 | tolerated | 0.14 | probably damaging | 0.984 |
| 42126938C>T | 1 | 1 | 0 | 2 | 0,1% | yes | -0.0489283204986387 | missense_variant | rs769157652 | tolerated | 0.22 | benign | 0.063 |
| 42126940delC | 1 | 0 | 0 | 1 | 0,1% | yes | -0.218086598672027 | frameshift_variant | | - | - | - | - |
| 42126944_42126945insT | 0 | 0 | 1 | 1 | 0,1% | no | | frameshift_variant | | - | - | - | - |
| 42126944C>T | 0 | 0 | 1 | 1 | 0,1% | no | | missense_variant | rs78762568 | tolerated | 0.22 | benign | 0 |
| 42126963C>T | 16 | 2 | 0 | 18 | 1,1% | yes | -0.00251656770706176 | synonymous_variant | rs28371732 | - | - | - | - |
| 42127001G>A | 353 | 126 | 51 | 530 | 33,2% | yes | -0.058850394544319715 | intron_variant | rs4987144 | - | - | - | - |
| 42127019G>T | 1 | 0 | 0 | 1 | 0,1% | no | | intron_variant | rs1008147170 | - | - | - | - |
| 42127173C>T | 1 | 0 | 0 | 1 | 0,1% | no | | intron_variant | rs1249410876 | - | - | - | - |
| 42127207C>T | 353 | 126 | 51 | 530 | 33,2% | yes | -0.026299823547962564 | intron_variant | | - | - | - | - |
| 42127209T>C | 250 | 61 | 30 | 341 | 21,4% | yes | -0.046035356570000484 | intron_variant | rs2004511 | - | - | - | - |
| 42127356G>T | 29 | 5 | 3 | 37 | 2,3% | yes | -0.036148767544496224 | intron_variant | rs28371729 | - | - | - | - |
| 42127385C>A | 1 | 0 | 0 | 1 | 0,1% | no | | intron_variant | rs891742278 | - | - | - | - |
| 42127398A>G | 5 | 9 | 4 | 18 | 1,1% | no | | intron_variant | rs28578778 | - | - | - | - |
| 42127407T>G | 603 | 184 | 82 | 869 | 54,4% | yes | -0.08095961630167535 | intron_variant | rs1985842 | - | - | - | - |
| 42127458G>A | 1 | 0 | 0 | 1 | 0,1% | yes | 0.174335300922393 | missense_variant | rs146271511 | tolerated | 0.13 | possibly damaging | 0.695 |
| 42127526_42127527insT | 0 | 1 | 0 | 1 | 0,1% | no | | frameshift_variant | | - | - | - | - |
| 42127526C>T | 2 | 1 | 0 | 3 | 0,2% | yes | 0.130049116392221 | missense_variant | rs1058172 | deleterious | 0.01 | probably damaging | 0.999 |
| 42127556T>C | 3 | 1 | 0 | 4 | 0,3% | yes | -0.08524450659751888 | missense_variant | rs202102799 | deleterious | 0 | probably damaging | 0.993 |
| 42127565T>C | 3 | 1 | 0 | 4 | 0,3% | yes | -0.165771216154098 | missense_variant | rs61736517 | tolerated | 1 | benign | 0 |
| 42127611C>T | 3 | 1 | 1 | 5 | 0,3% | yes | -0.16890266655038766 | missense_variant | rs78209835 | deleterious | 0.05 | benign | 0 |
| 42127631C>G | 1 | 2 | 0 | 3 | 0,2% | yes | -0.496762812137603 | missense_variant | rs141009491 | deleterious | 0 | probably damaging | 0.971 |
| 42127634C>A | 1 | 0 | 0 | 1 | 0,1% | yes | NA | missense_variant | rs3915951 | deleterious | 0.03 | benign | 0.037 |
| 42127644C>T | 2 | 0 | 0 | 2 | 0,1% | no | | intron_variant | rs376751897 | - | - | - | - |
| 42127677_42127687delGTCCGGCCCTG | 1 | 0 | 0 | 1 | 0,1% | no | | intron_variant | rs1237764592 | - | - | - | - |

| Variant (NC_000022.11:g.) | CYPTAM | BRUT | EIND | Total | Minor allele frequency (1596 alleles) | included neural network | Predicted contribution to allele activity based on neural network | consequence | rs-number | sift prediction | sift score | polyphen prediction | polyphen score |
|---|---|---|---|---|---|---|---|---|---|---|---|---|---|
| 42127694_42127695delCT | 1 | 0 | 0 | 1 | 0,1% | no | | intron_variant | | - | - | - | - |
| 42127698T>C | 1 | 0 | 0 | 1 | 0,1% | no | | intron_variant | | - | - | - | - |
| 42127700G>T | 1 | 0 | 0 | 1 | 0,1% | no | | intron_variant | | - | - | - | - |
| 42127707A>G | 1 | 0 | 0 | 1 | 0,1% | no | | intron_variant | | - | - | - | - |
| 42127718G>A | 1 | 0 | 0 | 1 | 0,1% | no | | intron_variant | rs1414906948 | - | - | - | - |
| 42127721C>T | 1 | 0 | 0 | 1 | 0,1% | no | | intron_variant | | - | - | - | - |
| 42127734T>G | 1 | 0 | 0 | 1 | 0,1% | no | | intron_variant | | - | - | - | - |
| 42127740T>C | 1 | 0 | 0 | 1 | 0,1% | no | | intron_variant | | - | - | - | - |
| 42127743A>C | 1 | 0 | 0 | 1 | 0,1% | no | | intron_variant | | - | - | - | - |
| 42127753T>G | 1 | 0 | 0 | 1 | 0,1% | no | | intron_variant | | - | - | - | - |
| 42127755G>A | 1 | 0 | 0 | 1 | 0,1% | no | | intron_variant | | - | - | - | - |
| 42127761C>T | 1 | 0 | 0 | 1 | 0,1% | yes | 0.0785754323005676 | intron_variant | rs267608291 | - | - | - | - |
| 42127774A>T | 1 | 0 | 0 | 1 | 0,1% | no | | intron_variant | rs1423323203 | - | - | - | - |
| 42127778T>C | 1 | 0 | 0 | 1 | 0,1% | no | | intron_variant | | - | - | - | - |
| 42127779T>C | 1 | 0 | 0 | 1 | 0,1% | no | | intron_variant | | - | - | - | - |
| 42127782T>G | 1 | 0 | 0 | 1 | 0,1% | no | | intron_variant | | - | - | - | - |
| 42127791G>A | 1 | 0 | 0 | 1 | 0,1% | no | | intron_variant | rs536709258 | - | - | - | - |
| 42127792C>G | 1 | 0 | 0 | 1 | 0,1% | no | | intron_variant | rs765006570 | - | - | - | - |
| 42127803C>T | 93 | 34 | 16 | 143 | 9,0% | yes | -0.37951637920601106 | intron_variant | rs28371725 | - | - | - | - |
| 42127811G>A | 4 | 0 | 2 | 6 | 0,4% | no | | intron_variant | rs143276168 | - | - | - | - |
| 42127813C>T | 1 | 0 | 0 | 1 | 0,1% | no | | intron_variant | | - | - | - | - |
| 42127820A>C | 1 | 0 | 0 | 1 | 0,1% | no | | intron_variant | | - | - | - | - |
| 42127821C>T | 1 | 0 | 0 | 1 | 0,1% | no | | intron_variant | rs1014423327 | - | - | - | - |
| 42127824T>G | 1 | 0 | 0 | 1 | 0,1% | no | | intron_variant | | - | - | - | - |
| 42127825T>G | 1 | 0 | 0 | 1 | 0,1% | no | | intron_variant | | - | - | - | - |
| 42127826T>C | 1 | 0 | 0 | 1 | 0,1% | no | | intron_variant | | - | - | - | - |
| 42127832A>C | 1 | 0 | 0 | 1 | 0,1% | no | | intron_variant | rs751061716 | - | - | - | - |
| 42127852C>T | 6 | 0 | 0 | 6 | 0,4% | yes | -0.46053318055295134 | synonymous_variant | rs79292917 | - | - | - | - |
| 42127853G>A | 1 | 0 | 0 | 1 | 0,1% | yes | NA | missense_variant | rs140513104 | deleterious | 0 | possibly damaging | 0.558 |
| 42127855A>G | 1 | 0 | 0 | 1 | 0,1% | no | | synonymous_variant | rs28371724 | - | - | - | - |
| 42127856T>G | 0 | 1 | 0 | 1 | 0,1% | no | | missense_variant | rs5030867 | deleterious | 0 | probably damaging | 1 |
| 42127940_42127941insA | 0 | 1 | 0 | 1 | 0,1% | no | | frameshift_variant | | - | - | - | - |
| 42127941G>A | 353 | 127 | 51 | 531 | 33,3% | yes | -0.05377423686868765 | missense_variant | rs16947 | tolerated | 0.21 | benign | 0.062 |
| 42127973T>C | 0 | 1 | 0 | 1 | 0,1% | no | | missense_variant | rs1135829 | tolerated | 0.08 | benign | 0.017 |
| 42127996G>A | 1 | 0 | 0 | 1 | 0,1% | no | | intron_variant | rs371181941 | - | - | - | - |
| 42128042A>G | 1 | 0 | 0 | 1 | 0,1% | no | | intron_variant | rs971733628 | - | - | - | - |
| 42128071C>G | 5 | 4 | 1 | 10 | 0,6% | no | | intron_variant | rs187203531 | - | - | - | - |
| 42128088G>A | 1 | 0 | 0 | 1 | 0,1% | no | | intron_variant | rs74516776 | - | - | - | - |
| 42128130C>T | 4 | 0 | 0 | 4 | 0,3% | yes | 0.022598231385327648 | intron_variant | rs28371721 | - | - | - | - |
| 42128136C>T | 0 | 1 | 0 | 1 | 0,1% | no | | intron_variant | rs372521768 | - | - | - | - |
| 42128176_42128178delTCT | 37 | 11 | 3 | 51 | 3,2% | yes | -0.37228316068649164 | inframe_deletion | rs5030656 | - | - | - | - |
| 42128176delT | 1 | 0 | 0 | 1 | 0,1% | yes | NA | frameshift_variant | rs28371720 | - | - | - | - |
| 42128216G>T | 2 | 0 | 1 | 3 | 0,2% | yes | -0.0146875381469726 | synonymous_variant | rs28371718 | - | - | - | - |
| 42128218delG | 27 | 0 | 0 | 27 | 1,7% | yes | 0.047781497582548245 | frameshift_variant | rs72549352 | - | - | - | - |
| 42128242delT | 22 | 4 | 7 | 33 | 2,1% | yes | -1 | frameshift_variant | rs35742686 | - | - | - | - |
| 42128251_42128254delTTAG | 1 | 0 | 1 | 2 | 0,1% | yes | -0.54435521364212 | frameshift_variant | rs72549353 | - | - | - | - |
| 42128308C>A | 13 | 1 | 2 | 16 | 1,0% | yes | 0.019396603107452302 | missense_variant | rs28371717 | tolerated | 0.41 | benign | 0.097 |
| 42128321A>G | 4 | 3 | 2 | 9 | 0,6% | yes | 0.228823016128404 | synonymous_variant | rs17002852 | - | - | - | - |
| 42128327G>T | 0 | 1 | 0 | 1 | 0,1% | no | | synonymous_variant | rs1462273327 | - | - | - | - |
| 42128438C>T | 2 | 1 | 1 | 4 | 0,3% | no | | intron_variant | rs79650744 | - | - | - | - |
| 42128500C>T | 13 | 0 | 0 | 13 | 0,8% | yes | -0.1153512736167856 | intron_variant | rs267608300 | - | - | - | - |
| 42128519G>A | 1 | 0 | 0 | 1 | 0,1% | no | | intron_variant | rs878981790 | - | - | - | - |
| 42128631C>T | 1 | 0 | 0 | 1 | 0,1% | no | | intron_variant | rs1468364907 | - | - | - | - |
| 42128693_42128694insC | 0 | 1 | 0 | 1 | 0,1% | no | | intron_variant | | - | - | - | - |
| 42128694T>C | 250 | 59 | 30 | 339 | 21,2% | yes | 0.05622132786746358 | intron_variant | rs2267447 | - | - | - | - |
| 42128736C>T | 1 | 0 | 0 | 1 | 0,1% | no | | intron_variant | rs761521669 | - | - | - | - |

| Variant (NC_000022.11:g.) | CYPTAM | BRUT | EIND | Total | Minor allele frequency (1596 alleles) | included neural network | Predicted contribution to allele activity based on neural network | consequence | rs-number | sift prediction | sift score | polyphen prediction | polyphen score |
|---|---|---|---|---|---|---|---|---|---|---|---|---|---|
| 42128741G>A | 1 | 0 | 1 | 2 | 0,1% | no | | intron_variant | rs113889384 | - | - | - | - |
| 42128793A>G | 6 | 1 | 0 | 7 | 0,4% | no | | synonymous_variant | rs28371713 | - | - | - | - |
| 42128815C>T | 15 | 6 | 2 | 23 | 1,4% | yes | -0.327434092760086 | missense_variant | rs5030866 | tolerated | 0.53 | benign | 0.189 |
| 42128858C>G | 0 | 1 | 0 | 1 | 0,1% | no | | missense_variant | | tolerated | 0.3 | benign | 0 |
| 42128922A>G | 6 | 0 | 0 | 6 | 0,4% | yes | 0.10502272844314499 | synonymous_variant | rs111606937 | - | - | - | - |
| 42128945C>T | 230 | 55 | 27 | 312 | 19,5% | yes | -0.22914504557519727 | splice_acceptor_variant | rs3892097 | - | - | - | - |
| 42128946_42128947insT | 0 | 1 | 0 | 1 | 0,1% | no | | splice_region_variant | | - | - | - | - |
| 42129084delA | 15 | 7 | 2 | 24 | 1,5% | yes | -0.616389781236648 | frameshift_variant | rs5030655 | - | - | - | - |
| 42129087G>C | 2 | 0 | 1 | 3 | 0,2% | yes | 0.2265237666148455 | missense_variant | rs78482768 | tolerated | 0.88 | benign | 0.003 |
| 42129130C>G | 603 | 186 | 79 | 868 | 54,4% | yes | -0.15171797035812631 | synonymous_variant | rs1058164 | - | - | - | - |
| 42129174C>A | 1 | 0 | 0 | 1 | 0,1% | yes | 0.237182915210723 | missense_variant | rs1135823 | tolerated | 0.39 | possibly damaging | 0.52 |
| 42129178G>A | 4 | 1 | 0 | 5 | 0,3% | no | | synonymous_variant | rs61736507 | - | - | - | - |
| 42129180A>T | 2 | 0 | 0 | 2 | 0,1% | yes | 0.5293659449718591 | missense_variant | rs1135822 | tolerated | 1 | benign | 0 |
| 42129184C>T | 0 | 1 | 0 | 1 | 0,1% | no | | splice_region_variant | rs267608304 | - | - | - | - |
| 42129225A>C | 0 | 0 | 1 | 1 | 0,1% | no | | intron_variant | rs370267861 | - | - | - | - |
| 42129228A>G | 0 | 0 | 1 | 1 | 0,1% | no | | intron_variant | rs373668214 | - | - | - | - |
| 42129303T>G | 2 | 0 | 0 | 2 | 0,1% | no | | intron_variant | rs142302759 | - | - | - | - |
| 42129319_42129320delTT | 1 | 0 | 0 | 1 | 0,1% | no | | intron_variant | rs1278131170 | - | - | - | - |
| 42129351A>G | 1 | 0 | 0 | 1 | 0,1% | no | | intron_variant | rs376056664 | - | - | - | - |
| 42129388G>A | 6 | 1 | 0 | 7 | 0,4% | no | | intron_variant | rs557722765 | - | - | - | - |
| 42129414G>C | 4 | 1 | 0 | 5 | 0,3% | no | | intron_variant | rs143170489 | - | - | - | - |
| 42129436A>G | 2 | 0 | 0 | 2 | 0,1% | no | | intron_variant | rs184086520 | - | - | - | - |
| 42129450delG | 1 | 0 | 0 | 1 | 0,1% | no | | intron_variant | | - | - | - | - |
| 42129461G>A | 1 | 0 | 0 | 1 | 0,1% | no | | intron_variant | | - | - | - | - |
| 42129498C>T | 1 | 0 | 0 | 1 | 0,1% | no | | intron_variant | rs189736703 | - | - | - | - |
| 42129545G>A | 16 | 6 | 2 | 24 | 1,5% | no | | intron_variant | rs267608277 | - | - | - | - |
| 42129581T>G | 1 | 0 | 0 | 1 | 0,1% | no | | intron_variant | rs1170216892 | - | - | - | - |
| 42129587C>T | 1 | 0 | 0 | 1 | 0,1% | no | | intron_variant | rs1183013541 | - | - | - | - |
| 42129614C>G | 2 | 0 | 0 | 2 | 0,1% | no | | intron_variant | rs180847475 | - | - | - | - |
| 42129623C>T | 80 | 19 | 10 | 109 | 6,8% | no | | intron_variant | rs1081004 | - | - | - | - |
| 42129643G>C | 2 | 0 | 0 | 2 | 0,1% | no | | intron_variant | rs186133763 | - | - | - | - |
| 42129722C>T | 1 | 0 | 0 | 1 | 0,1% | no | | intron_variant | rs368389952 | - | - | - | - |
| 42129726A>C | 1 | 0 | 0 | 1 | 0,1% | no | | intron_variant | rs78854695 | - | - | - | - |
| 42129731T>C | 1 | 0 | 0 | 1 | 0,1% | yes | -0.243858930342826 | splice_region_variant | rs267608289 | - | - | - | - |
| 42129754G>A | 23 | 13 | 6 | 42 | 2,6% | yes | 0.058333239961291326 | synonymous_variant | rs1081003 | - | - | - | - |
| 42129770G>A | 2 | 0 | 0 | 2 | 0,1% | yes | 0.14140502977480102 | missense_variant | rs28371706 | tolerated | 0.21 | benign | 0 |
| 42129793G>A | 0 | 0 | 1 | 1 | 0,1% | no | | synonymous_variant | rs200269944 | - | - | - | - |
| 42129795_42129796insC | 0 | 1 | 0 | 1 | 0,1% | no | | frameshift_variant | | - | - | - | - |
| 42129796G>C | 225 | 46 | 23 | 294 | 18,4% | yes | 0.1486052139837259 | synonymous_variant | rs28371705 | - | - | - | - |
| 42129809T>C | 225 | 48 | 24 | 297 | 18,6% | yes | -0.25058796488421997 | missense_variant | rs28371704 | tolerated | 1 | benign | 0 |
| 42129819G>T | 225 | 48 | 23 | 296 | 18,5% | yes | -0.036173526027646376 | missense_variant | rs28371703 | deleterious | 0.02 | possibly damaging | 0.927 |
| 42129950A>C | 610 | 188 | 80 | 878 | 55,0% | yes | -0.29604015773312636 | intron_variant | rs28371702 | - | - | - | - |
| 42129950delA | 1 | 0 | 0 | 1 | 0,1% | no | | intron_variant | | - | - | - | - |
| 42130047G>C | 578 | 174 | 74 | 826 | 51,8% | yes | 0.04334876873857956 | intron_variant | rs28371701 | - | - | - | - |
| 42130180C>T | 1 | 0 | 0 | 1 | 0,1% | no | | intron_variant | rs896350438 | - | - | - | - |
| 42130343T>G | 0 | 1 | 0 | 1 | 0,1% | no | | intron_variant | rs867984132 | - | - | - | - |
| 42130469C>T | 1 | 0 | 1 | 2 | 0,1% | no | | intron_variant | rs35970455 | - | - | - | - |
| 42130475T>C | 1 | 0 | 1 | 2 | 0,1% | no | | intron_variant | rs34291018 | - | - | - | - |
| 42130482C>A | 603 | 188 | 79 | 870 | 54,5% | yes | 0.10251049786610479 | intron_variant | rs28371699 | - | - | - | - |
| 42130522G>A | 44 | 15 | 7 | 66 | 4,1% | no | | intron_variant | rs29001678 | - | - | - | - |
| 42130538C>A | 0 | 0 | 1 | 1 | 0,1% | no | | intron_variant | rs1255242741 | - | - | - | - |
| 42130547T>C | 354 | 127 | 52 | 533 | 33,4% | yes | 0.020412143103732624 | intron_variant | | - | - | - | - |
| 42130552G>A | 1 | 0 | 0 | 1 | 0,1% | no | | intron_variant | | - | - | - | - |
| 42130559T>G | 354 | 127 | 52 | 533 | 33,4% | yes | -0.016224358148486787 | intron_variant | rs28695233 | - | - | - | - |
| 42130560C>G | 354 | 127 | 52 | 533 | 33,4% | yes | -0.02308920692216708 | intron_variant | rs29001518 | - | - | - | - |
| 42130565A>G | 354 | 127 | 52 | 533 | 33,4% | yes | 0.06461922083170608 | intron_variant | rs1080998 | - | - | - | - |

| Variant (NC_000022.11:g.) | CYPTAM | BRUT | EIND | Total | Minor allele frequency (1596 alleles) | included neural network | Predicted contribution to allele activity based on neural network | consequence | rs-number | sift prediction | sift score | polyphen prediction | polyphen score |
|---|---|---|---|---|---|---|---|---|---|---|---|---|---|
| 42130569G>C | 354 | 127 | 52 | 533 | 33,4% | yes | -0.24267589335388073 | intron_variant | rs1080997 | - | - | - | - |
| 42130571G>T | 354 | 127 | 52 | 533 | 33,4% | yes | 0.1175802367055315 | intron_variant | rs1080996 | - | - | - | - |
| 42130578C>G | 354 | 127 | 52 | 533 | 33,4% | yes | -0.005272804071269889 | intron_variant | rs1080995 | - | - | - | - |
| 42130594_42130595insT | 0 | 2 | 0 | 2 | 0,1% | no | | intron_variant | | - | - | - | - |
| 42130655_42130656insA | 0 | 0 | 2 | 2 | 0,1% | no | | frameshift_variant | rs774671100 | - | - | - | - |
| 42130657G>A | 1 | 0 | 0 | 1 | 0,1% | no | | synonymous_variant | rs757862366 | - | - | - | - |
| 42130667C>T | 1 | 0 | 0 | 1 | 0,1% | yes | -0.521055400371551 | missense_variant | rs118203758 | deleterious | 0.02 | probably damaging | 0.953 |
| 42130692G>A | 248 | 60 | 29 | 337 | 21,1% | yes | -0.07567874934073372 | missense_variant | rs1065852 | deleterious | 0.02 | possibly damaging | 0.687 |
| 42130710G>A | 5 | 1 | 0 | 6 | 0,4% | yes | -0.103839337825775 | missense_variant | rs138100349 | tolerated | 0.07 | benign | 0.021 |
| 42130761_42130762insT | 0 | 0 | 2 | 2 | 0,1% | no | | frameshift_variant | | - | - | - | - |
| 42130761C>T | 61 | 28 | 8 | 97 | 6,1% | yes | -0.027715351754274285 | missense_variant | rs769258 | tolerated | 0.14 | benign | 0.024 |
| 42130773C>T | 2 | 0 | 1 | 3 | 0,2% | yes | -0.09531869985836064 | missense_variant | rs72549358 | tolerated | 0.22 | benign | 0 |
| 42130929T>C | 1 | 0 | 0 | 1 | 0,1% | yes | 0.10511073372503 | upstream_gene_variant | rs267608272 | - | - | - | - |
| 42131000G>A | 0 | 1 | 0 | 1 | 0,1% | no | | upstream_gene_variant | rs912915978 | - | - | - | - |
| 42131062T>A | 1 | 0 | 0 | 1 | 0,1% | no | | upstream_gene_variant | rs1346206732 | - | - | - | - |
| 42131114A>G | 5 | 1 | 0 | 6 | 0,4% | no | | upstream_gene_variant | rs572914357 | - | - | - | - |
| 42131156C>T | 18 | 8 | 0 | 26 | 1,6% | no | | upstream_gene_variant | rs1080992 | - | - | - | - |
| 42131222G>A | 1 | 1 | 0 | 2 | 0,1% | yes | -0.0395302993694236 | upstream_gene_variant | rs566383351 | - | - | - | - |
| 42131379C>T | 11 | 0 | 1 | 12 | 0,8% | no | | upstream_gene_variant | rs769257 | - | - | - | - |
| 42131469C>T | 350 | 127 | 51 | 528 | 33,1% | yes | -0.02665604532520481 | upstream_gene_variant | rs28633410 | - | - | - | - |
| 42131493C>T | 0 | 0 | 1 | 1 | 0,1% | no | | upstream_gene_variant | rs958348536 | - | - | - | - |
| 42131531G>A | 354 | 127 | 51 | 532 | 33,3% | yes | -0.013184694660057422 | upstream_gene_variant | rs28624811 | - | - | - | - |
| 42131546_42131547delTC | 1 | 0 | 0 | 1 | 0,1% | yes | -0.0190738309697214 | upstream_gene_variant | rs536645539 | - | - | - | - |
| 42131610G>C | 2 | 0 | 0 | 2 | 0,1% | no | | upstream_gene_variant | rs930275829 | - | - | - | - |
| 42131631A>T | 3 | 1 | 0 | 4 | 0,3% | no | | upstream_gene_variant | rs113127784 | - | - | - | - |
| 42131681C>T | 0 | 1 | 0 | 1 | 0,1% | no | | upstream_gene_variant | rs566307819 | - | - | - | - |
| 42131775C>T | 0 | 1 | 1 | 2 | 0,1% | no | | upstream_gene_variant | rs567431353 | - | - | - | - |
| 42131791C>T | 248 | 60 | 29 | 337 | 21,1% | yes | -0.22362225330200997 | upstream_gene_variant | rs1080989 | - | - | - | - |
| 42131894delT | 898 | 2 | 4 | 904 | 56,6% | no | | upstream_gene_variant | rs375413467 | - | - | - | - |
| 42131908C>T | 1 | 0 | 0 | 1 | 0,1% | no | | upstream_gene_variant | rs1250789569 | - | - | - | - |
| 42131914_42131915delCT | 0 | 0 | 1 | 1 | 0,1% | no | | upstream_gene_variant | | - | - | - | - |
| 42131915T>G | 1 | 0 | 0 | 1 | 0,1% | no | | upstream_gene_variant | rs62625688 | - | - | - | - |
| 42131927C>A | 3 | 1 | 0 | 4 | 0,3% | no | | upstream_gene_variant | rs528127834 | - | - | - | - |
| 42131990delC | 15 | 1 | 1 | 17 | 1,1% | no | | upstream_gene_variant | rs757622584 | - | - | - | - |
| 42132026T>C | 604 | 187 | 80 | 871 | 54,6% | yes | 0.30988314411907114 | upstream_gene_variant | rs28735595 | - | - | - | - |
| 42132027_42132028insC | 0 | 1 | 0 | 1 | 0,1% | no | | upstream_gene_variant | rs1080988 | - | - | - | - |
| 42132045_42132049delTTTTT | 1 | 0 | 0 | 1 | 0,1% | no | | upstream_gene_variant | rs267608321 | - | - | - | - |
| 42132046_42132049delTTTT | 3 | 0 | 0 | 3 | 0,2% | no | | upstream_gene_variant | rs267608321 | - | - | - | - |
| 42132047_42132049delTTT | 6 | 0 | 0 | 6 | 0,4% | no | | upstream_gene_variant | rs267608321 | - | - | - | - |
| 42132048_42132049delTT | 244 | 40 | 6 | 290 | 18,2% | no | | upstream_gene_variant | rs267608321 | - | - | - | - |
| 42132049_42132050insT | 109 | 34 | 19 | 162 | 10,2% | no | | upstream_gene_variant | rs267608321 | - | - | - | - |
| 42132049_42132050insTT | 194 | 55 | 17 | 266 | 16,7% | no | | upstream_gene_variant | rs267608321 | - | - | - | - |
| 42132049_42132050insTTT | 86 | 57 | 28 | 171 | 10,7% | no | | upstream_gene_variant | rs267608321 | - | - | - | - |
| 42132049_42132050insTTTT | 0 | 13 | 5 | 18 | 1,1% | no | | upstream_gene_variant | rs267608321 | - | - | - | - |
| 42132049delT | 259 | 79 | 33 | 371 | 23,2% | no | | upstream_gene_variant | rs267608321 | - | - | - | - |
| 42132138C>G | 0 | 0 | 1 | 1 | 0,1% | no | | upstream_gene_variant | rs757984982 | - | - | - | - |
| 42132217G>A | 250 | 62 | 30 | 342 | 21,4% | yes | -0.189309422790393 | upstream_gene_variant | rs28588594 | - | - | - | - |
| 42132375G>C | 253 | 93 | 34 | 380 | 23,8% | yes | 0.06632569069120157 | upstream_gene_variant | rs1080985 | - | - | - | - |
| 42132377delG | 1 | 0 | 0 | 1 | 0,1% | no | | upstream_gene_variant | | - | - | - | - |
| 42132411C>T | 1 | 0 | 0 | 1 | 0,1% | no | | upstream_gene_variant | | - | - | - | - |
| 42132561_42132562insT | 0 | 1 | 0 | 1 | 0,1% | no | | upstream_gene_variant | | - | - | - | - |
| 42132561C>T | 349 | 126 | 50 | 525 | 32,9% | yes | 0.06237363234918848 | upstream_gene_variant | rs1080983 | - | - | - | - |
| 42132577G>C | 0 | 1 | 0 | 1 | 0,1% | no | | upstream_gene_variant | rs1374920910 | - | - | - | - |
| 42132589C>T | 0 | 1 | 0 | 1 | 0,1% | no | | upstream_gene_variant | rs936292274 | - | - | - | - |
| 42132590G>A | 0 | 1 | 0 | 1 | 0,1% | no | | upstream_gene_variant | rs1054718426 | - | - | - | - |
| Del | 39 | 10 | 8 | 57 | 3,6% | yes | -1 | | | | | | |
| Dup | 13 | 4 | 3 | 20 | 1,3% | yes | 0.2267 | | | | | | |
